# EWI-2 Inhibits Cell-Cell Fusion at the HIV-1 Virological Presynapse

**DOI:** 10.1101/296251

**Authors:** Emily E. Whitaker, Nicholas J. Matheson, Sarah Perlee, Phillip B. Munson, Menelaos Symeonides, Markus Thali

**Affiliations:** University of Vermont, Department of Microbiology and Molecular Genetics, Burlington, VT, USA; University of Vermont, Graduate Program in Cellular, Molecular, and Biomedical Sciences, Burlington, VT, USA; Department of Medicine, University of Cambridge, Cambridge, UK; Cambridge Institute for Therapeutic Immunology and Infectious Disease (CITIID), University of Cambridge, Cambridge, UK; University of Vermont, Department of Pathology and Laboratory Medicine, Burlington, VT, USA

**Keywords:** EWI-2, IGSF8, tetraspanin, HIV, cell-cell fusion, virological synapse, T cell, syncytia

## Abstract

Cell-to-cell transfer of virus particles at the Env-dependent virological synapse (VS) is a highly efficient mode of HIV-1 transmission. While cell-cell fusion could be triggered at the VS, leading to the formation of syncytia and preventing exponential growth of the infected cell population, this is strongly inhibited by both viral (Gag) and host (ezrin and tetraspanins) proteins. Here, we identify EWI-2, a protein that was previously shown to associate with ezrin and tetraspanins, as a host factor that contributes to the inhibition of Env-mediated cell-cell fusion. Using quantitative fluorescence microscopy, shRNA knockdowns, and cell-cell fusion assays, we show that EWI-2 accumulates at the presynaptic terminal (i.e. the producer cell side of the VS), where it contributes to the fusion-preventing activities of the other viral and cellular components. We also find that EWI-2, like tetraspanins, is downregulated upon HIV-1 infection, mostly by Vpu. Despite strong inhibition of fusion at the VS, T cell-based syncytia do form *in vivo* and in physiologically relevant culture systems, but they remain small. In regard to that, we demonstrate that EWI-2 and CD81 levels are restored on the surface of syncytia, where they (presumably) continue to act as fusion inhibitors. This study documents a new role for EWI-2 as an inhibitor of HIV-1-induced cell-cell fusion, and provides novel insight into how syncytia are prevented from fusing indefinitely.

## 1. Introduction

HIV-1 spreads between T cells primarily through two modes of transmission: the release of cell-free virus particles followed by their uptake by (more or less distantly located) cells expressing the viral receptor/co-receptor, and cell-to-cell transmission of particles to an adjacent cell via the virological synapse (VS), i.e. when infected and uninfected cells transiently align. Formation of the HIV-1 VS is initiated by the viral envelope glycoprotein (Env) on the surface of productively infected cells binding to its receptor, CD4, on target T cells [1], and is followed by polarization of Gag at the cell-cell contact site [1,2]. Virus particles are then released in high concentrations towards the target cell [3], facilitating efficient infection while also possibly shielding virus particles from some neutralizing antibodies ([4] and recently reviewed in [5]). Indeed, as demonstrated in a recent study using physiologically relevant cell culture systems [6], it is possible that virus that is not released in close proximity to a target cell is rapidly inactivated, emphasizing the importance of VS-mediated transmission. Given, however, that Env is fusogenic at neutral pH, it would seem likely at first that VS-mediated contacts should frequently result in cell-cell fusion, thus forming a multinucleated infected cell (syncytium). While we now know that small, T cell-based syncytia arise early in HIV-1 infection and can spread virus by cell-cell contact [7–12], the majority of infected T cells observed in lymphoid tissue are mononucleated, documenting that most HIV-1 VSs ultimately result in complete cell separation and generation of a new, productively infected cell. This is likely due to tight regulation at the VS that acts to prevent excessive syncytium formation (reviewed in [13,14]).

Multiple independent studies have identified viral and host functions which, together, prevent excessive HIV-1-induced cell-cell fusion at the VS. Firstly, Env is rapidly downregulated from the surface of infected cells in the absence of Gag [15,16]. Secondly, upon Gag multimerization at the plasma membrane, Env is trapped by immature Gag through Env’s cytoplasmic tail and maintained in a poorly fusogenic state [17]. This trapping by Gag ends only after Env’s incorporation into virus particles, when Gag precursor gets cleaved, i.e. upon maturation [18–21]. The residual fusion activity of Gag-trapped Env on infected cells has been shown to be inhibited by several host membrane proteins that accumulate at the producer cell side of the VS, including tetraspanins and phosphorylated ezrin (p-ezrin) [22–24]. Tetraspanins inhibit HIV-1-induced cell-cell fusion at a post-hemifusion stage [23], while ezrin is implicated in F-actin organization and recruitment of the tetraspanin CD81 to the VS [24]. It remains unclear how and whether these protein functions are coordinated, though based on other cell-cell fusion regulation paradigms (discussed below), additional host proteins are likely required to mediate efficient inhibition of HIV-1-induced fusion by tetraspanins and ezrin.

EWI-F (CD9P-1/FPRP) is an immunoglobulin superfamily (IgSF) member and partner of tetraspanins CD9 and CD81 [25]. EWI-F was shown to be a potent inhibitor of cell-cell fusion in myoblasts, where EWI-F knockout resulted in more frequent fusion than CD9/CD81 double knockout [26]. However, EWI-F is poorly expressed in T cells [27], the primary host cell type for HIV-1. A related protein, EWI-2 (IGSF8/PGRL) [28,29], which also associates with tetraspanins and is expressed in T cells [25,27], has been documented to play a role in HCV entry [30,31] and T cell immunological synapse (IS) formation [32]. That latter study also suggested that EWI-2 has a yet undetermined involvement in HIV-1 particle production [32]. Furthermore, both EWI-F and EWI-2 interact with ezrin to organize the cytoskeleton in concert with tetraspanins [27]. EWI-2 thus lies at the nexus of tetraspanins, ezrin, and the actin cytoskeleton (which can also inhibit cell-cell fusion; [33]).

## 2. Materials and Methods

### 2.1 Cell Lines and Cell Culture

The following cells were obtained through the NIH AIDS Reagent Program, Division of AIDS, NIAID, NIH: HeLa cells from Dr. Richard Axel [34], TZM-bl cells from Dr. John C. Kappes, Dr. Xiaoyun Wu, and Tranzyme Inc. [35–39], CEM.NKR CCR5+Luc+ (CEM-luc) cells from Dr. John Moore and Dr. Catherine Spenlehauer [40,41], CEM-T4 cells from Dr. J.P. Jacobs [42], and CEM-SS cells from Dr. Peter L. Nara [34,43,44].

HEK 293T, HeLa, and TZM-bl cells were maintained in Dulbecco’s Modification of Eagle’s Medium (DMEM) (Corning, Corning, NY, Cat. #10-017-CV) containing 10% fetal bovine serum (FBS; Corning, Cat. #35-010-CV) and antibiotics (100 units/mL penicillin and 100 μg/mL streptomycin; Invitrogen). CEM-luc cells were maintained in RPMI 1640 medium (Corning, Cat. #10-104-CV) supplemented with 10% FBS and 0.8 mg/mL geneticin sulfate (G418). CEM2n, a kind gift from R. Harris [45], and CEM-SS cells were maintained in RPMI medium supplemented with 10% FBS and antibiotics.

Human primary blood mononuclear cells (PBMCs) were isolated as buffy coats from whole blood of healthy donors by Ficoll density centrifugation. CD4^+^ T cells were enriched from PBMCs by negative selection using the MACS CD4^+^ T Cell Isolation Kit (Miltenyi Biotec, Auburn, CA, Cat. #130-096-533) or the EasySep Human CD4^+^ T Cell Isolation Kit (STEMCELL Technologies, Vancouver, BC, Canada, Cat. #17952) according to manufacturer’s instructions. Primary CD4^+^ T cells were activated in RPMI containing 10% FBS, 50 units/mL IL-2, antibiotics, and 5 μg/mL phytohemagglutinin. After 48 h of activation, cells were washed and subsequently maintained and expanded in the same medium but without phytohemagglutinin. Cells were used for infections at 4 to 7 days post isolation.

### 2.2 Antibodies

Mouse monoclonal antibody (mAb) to EWI-2 (8A12) was a kind gift from Dr. Eric Rubinstein [25]. Mouse mAb to HIV-1 p24 (AG3.0) was obtained through the NIH AIDS Reagent Program, Division of AIDS, NIAID, NIH, from Dr. Jonathan Allan [46]. Rabbit antiserum to HIV-1 p6 was a kind gift from David E. Ott. Rabbit polyclonal antibody (pAb) to HIV-1 p24 was obtained from Advanced Biotechnologies (Cat. #13-203-000). Secondary antibodies were as follows: Alexa Fluor 488-conjugated donkey pAb to mouse IgG (Invitrogen, Cat. #A21202), Alexa Fluor 488-conjugated donkey pAb to rabbit IgG (Invitrogen Cat. #A21206), Alexa Fluor 594-conjugated donkey pAb to mouse IgG (Invitrogen Cat. #R37115), Alexa Fluor 594-conjugated donkey pAb to rabbit IgG (Invitrogen Cat. #A21207), Alexa Fluor 647-conjugated donkey pAb to mouse IgG (Invitrogen Cat. #A31571), and Alexa Fluor 647-conjugated goat pAb to mouse IgG (Invitrogen Cat. #A21235). Zenon labeling of primary antibodies with either Alexa Fluor 488 or Alexa Fluor 594 was carried out using Zenon Labeling Kits according to the manufacturer’s instructions (Molecular Probes, Eugene, OR, Cat. #Z25002 and #Z25007).

### 2.3 Plasmids and Virus Strains

pcDNA3, pCDNA3.1, and pCMV SPORT6 (Invitrogen) were vectors for EWI-2, CD81, and L6 overexpression, respectively (EWI-2 was a kind gift from Dr. Eric Rubinstein; Université Paris-Sud, Villejuif, France). Proviral plasmids pNL4-3 and pNL4-3 ΔEnv (KFS) were kind gifts from Dr. Eric Freed (National Cancer Institute, Frederick, MD, USA) [47]. NL4-3-derived fluorescent protein-tagged proviral plasmids pNL-sfGI, pNL-sfGI ΔEnv, pNL-CI, and pNL-CI ΔEnv [10] were kind gifts from Dr. Benjamin Chen (Mount Sinai School of Medicine, New York, NY). Vesicular stomatitis virus glycoprotein (VSV-G) was used to pseudotype viral stocks produced in HEK 293T cells. The lentiviral vector FG12 [48], previously modified to include a puromycin resistance cassette [24], was further modified to remove the GFP reporter cassette by digestion with AfeI and PshAI and subsequent blunt-end religation.

### 2.4 Virus Stocks and Infections

VSV-G-pseudotyped virus stocks of NL4-3, NL4-3 ∆Env, NL-sfGI, NL-CI, and NL-CI ∆Env were produced in HEK 293T cells transfected with the proviral plasmid and pVSV-G (at 17:3 ratio) using calcium phosphate precipitation. For shRNA encoding lentiviruses, shEWI-2 and shScramble, stocks were produced in HEK 293T cells transfected with FG12-shRNA vector, ΔR8.2 packaging vector, and pVSV-G (at a ratio of 3:7:1. Supernatants were harvested 2 days after transfection, cleared by centrifugation at 2000 rcf for 10 min, filtered, and stored at −80 °C.

To infect CEM2n cells by spinoculation, two million cells were incubated with RPMI/10% FBS containing 90 μL of virus stock (resulting in ~3% of the cells being infected) or medium alone (for uninfected controls), for 20 min at 37 °C, followed by centrifugation at 1200 rcf for 2 h at 37 °C. Cell pellets were allowed to recover at 37 °C for 15 min, centrifuged at 300 rcf for 2 min, and resuspended in fresh RPMI/10% FBS. Cells were incubated at 37 °C, the medium was refreshed 2 days post infection, and the cells were used 1 day later for all subsequent experiments.

To infect primary CD4^+^ T cells, 1 or 2 million cells were incubated in RPMI/10% FBS/IL-2 containing 200 or 400 μL of virus, respectively, and spinoculated as described above. Cells were resuspended in fresh RPMI/10% FBS/PS/IL-2 and incubated at 37 °C/5% CO2. Cells were used 2 days post infection for all subsequent experiments.

To infect CEM-SS cells by shaking, one or two million cells suspended in CO_2_-independent medium (Gibco) supplemented with 10% FBS were mixed with VSV-G-pseudotyped virus stocks and shaken at 220 rpm for 2 h at 37 °C. Cells were then washed and plated in fresh RPMI/10% FBS, and used for experiments as described. For CEM-SS infection by spinoculation, the procedure was performed as described above with some modifications; one or two million cells were incubated in RPMI/10% FBS containing 40-50 μL (analyzing surface expression and post-synapse enrichment, respectively) of virus stock or medium alone (for uninfected controls). Following spinoculation, cells were incubated at 37 °C for 2 days before being used for subsequent experiments.

### 2.5 Imaging and quantification of EWI-2 accumulation at the VS

CEM-SS and primary CD4^+^ T cells were infected by shaking or spinoculation, respectively, with VSV-G-pseudotyped WT or ΔEnv virus then treated as follows: For CEM-SS cells, two days post infection, uninfected CEM-SS target cells were labeled with CMAC (Invitrogen) according to manufacturer’s instructions, mixed with infected cells at a 1:1 or 1:2 ratio (infected:target), seeded onto the microwell of a 35 mm glass-bottom dish (MatTek Corporation, Ashland, MA, Cat. #P35G-1.5-14-C) coated with poly-L-Lysine (Sigma), and incubated at 37° C for 3 to 4.5 h. Cells were then chilled on ice and surface-labeled with 1:200 mouse anti-EWI-2 mAb in RPMI/10% FBS for 45 min at 4 °C. Surface-labeled cells were fixed with 4% PFA in PBS at 4 °C for 10 min, and blocked and permeabilized overnight with 1% BSA and 0.2% Triton X-100 in PBS (block/perm buffer). All CEM-SS conditions were labeled with Alexa Fluor 647-conjugated anti-mouse secondary pAb in block/perm buffer at 1:500 dilution. Cells were subsequently stained with Alexa Fluor 594 Zenon-labeled anti-p24 AG3.0 mouse mAb, and fixed again with 4% PFA in PBS. Cells were kept in PBS for imaging.

For primary cells, uninfected cells were mixed with infected cells at a 1:1 ratio (infected:target), seeded onto 8-well glass-bottom plates (CellVis, Mountain View, CA, Cat. #C8-1.5H-N) coated with 1:10 poly-L-Lysine in double-distilled water (ddH_2_O), and incubated for 2 to 2.5 h at 37° C. Cells were surface-labeled for EWI-2 and fixed as above, then blocked and permeabilized with block/perm buffer for 10 min. Cells were then labeled with a mixture of rabbit anti-p24 and anti-p6 antibodies, each at 1:1000 dilution, in PBS with 1% BSA (block) for 45 min. Subsequently, cells were labeled with Alexa Fluor-conjugated secondary pAbs as indicated. Cells were kept in PBS for imaging.

To visualize only producer cell-associated EWI-2 at the VS, 10,000 target TZM-bl cells (which have nearly-undetectable levels of EWI-2; unpublished observation) were seeded onto 8-well glass-bottom plates coated with 1:10 poly-L-Lysine in ddH_2_O. The next day, those TZM-bl cells were labeled with CMAC at 1:250 dilution in serum-free DMEM, and then co-cultured with 150,000 CEM-SS cells (either uninfected or infected with NL-CI or NL-CI ∆Env 2 days prior as described above) per well for 2.5 hours at 37 °C in RPMI/10% FBS. The cells were then surface-labeled with 1:200 mouse anti-EWI-2 mAb in RPMI/10% FBS on ice for 45 min. Cells were subsequently fixed with 4% PFA in PBS and permeabilized with block/perm for 10 min. After permeabilization, the cells were labeled using a mixture of rabbit anti-p24 and anti-p6 antibodies, each at 1:1000 dilution, in block for 45 min. Cells were subsequently labeled using Alexa Fluor-conjugated secondary pAbs (anti-mouse-Alexa Fluor 647 and anti-rabbit-Alexa Fluor 488) each at 1:500 in block for 45 minutes. Cells were kept in PBS for imaging.

To visualize only target cell-associated EWI-2 at the VS, HeLa producer cells (which have nearly-undetectable levels of EWI-2; unpublished observation) were plated (10,000 cells per well) in 8-well glass-bottom plates coated with 1:10 poly-L-Lysine in ddH_2_O. Twenty-four hours later, cells were transfected with NL-sfGI, NL-sfGI ∆Env, or empty vector, using FuGENE6 transfection reagent at a ratio of 3:1 (FuGENE6:DNA) according to manufacturer’s instructions (Promega, Madison, WI, Cat. #E2691). Twenty-four hours post-transfection, 100,000-150,000 uninfected CEM-SS cells (labeled with CMAC at a 1:250 dilution in serum-free RPMI) were added to form VSs with provirus-transfected HeLa cells. After 2-2.5 h of coculture, cells were surface-labeled with 1:200 mouse anti-EWI-2 mAb in RPMI/10% FBS for 45 min at 4 °C. Surface-labeled cells were fixed with 4% PFA in PBS at 4 °C for 10 min, and then incubated with block/perm for 10 min, before labeling with a mixture of rabbit anti-p24 and anti-p6 antibodies, each at 1:1000 dilution, in block for 45 min. Subsequently, cells were labeled with secondary pAbs (anti-mouse-Alexa Fluor 647 and anti-rabbit-Alexa Fluor 594), each at 1:500 in block. Cells were kept in PBS for imaging.

Images were acquired on a DeltaVision epifluorescence microscope (GE/Applied Precision, Issaquah, WA, USA) with an Olympus IX-70 base using an Olympus 60× PlanApo 1.42 NA objective and equipped with a CoolSNAP HQ CCD camera (Photometrics). Images were imported into Fiji Version 2.0.0-rc-69/1.52p [49] for analysis following deconvolution and cropping using Softworx software. The VS was identified using the Gag channel and the level of EWI-2 accumulation was determined by measuring its signal intensity at the VS. For ΔEnv controls, cell-cell contacts were identified using the differential interference contrast (DIC) channel and treated analogous to a VS. The EWI-2-associated signal intensity at non-contact sites was determined by manually outlining the surface of the cell, excluding any regions that were in contact with an adjacent cell, and calculating the mean EWI-2 intensity within the selected area. To determine the level of enrichment at the VS (or cell-cell contact for ΔEnv controls), an “unbiased” approach was applied to account for EWI-2 signal contributed by both the target and producer cell at each VS/contact. Enrichment was calculated as the EWI-2 signal intensity at the VS/contact divided by the sum of the EWI-2 signal at non-contact sites of the producer and target cell in that particular VS/contact. A “biased” approach, where only the producer cell’s non-contact sites were used to normalize the VS/contact signal, yielded very similar results to the unbiased approach described above (unpublished observations).

### 2.6 Proteomic analysis of EWI-2 levels in HIV-1 infected cells

To identify HIV-1-dependent changes in abundance of total EWI-2, we re-analysed data from 2 previous studies [50,51]. In brief, primary human CD4^+^ T cells were infected with pNL4-3-∆Env-Nef-P2A-SBP-∆LNGFR (HIV-AFMACS) at MOI≤0.5, enriched by Antibody-Free Magnetic Cell Sorting (AFMACS) [52] and analysed 48 h after infection [51]. CEM-T4 T cells were infected with pNL4-3-∆Env-EGFP at MOI=1.5 and analysed 48 h after infection [50]. TMT-labeled tryptic peptides from whole cell lysates were subjected to off-line High pH Reversed-Phase (HpRP)-HPLC fractionation and analysed using an Orbitrap Fusion Tribrid mass spectrometer (Thermo Scientific) coupled to a Dionex UltiMate 3000 UHPLC (Thermo Scientific). Details of sample processing and data analysis have been previously described [50,51] and proteomic data from primary human CD4^+^ T cells are available from the ProteomeX-change Consortium using dataset identifier PXD012263 (http://proteomecentral.proteomexchange.org).

To characterise HIV-1-dependent changes in abundance of plasma membrane EWI-2, we re-analysed data from a previous study [53]. In brief, for the TMT-based time course experiment, CEM-T4 T cells were infected with pNL4-3-∆Env-EGFP at MOI=10 and analysed at the indicated time points after infection. For the SILAC-based single time point experiments, cells were pre-labeled with light, medium or heavy lysine and arginine and either infected with WT or Vpu-/Nef-deficient pNL4-3-∆Env-EGFP at MOI=10 and analysed 72 h after infection, or transduced with GFP or Vpu/Nef and selected with puromycin. Sialylated cell surface glycoproteins were enriched by selective aminooxy-biotinylation followed by immunoaffinity purification using streptavidin-conjugated beads (Plasma Membrane Profiling). Tryptic peptides were labeled with TMT reagents (time course experiment only), subjected to off-line High pH Reversed-Phase (HpRP)-HPLC fractionation and analysed using an Orbitrap Fusion Tribrid mass spectrometer (Thermo Scientific) coupled to a Dionex UltiMate 3000 UHPLC (Thermo Scientific). Details of sample processing and data analysis have been previously described [53] and time course proteomic data are available from the ProteomeX-change Consortium using dataset identifier PXD002934 (http://proteomecentral.proteomexchange.org).

### 2.7 Determining surface levels of EWI-2 by microscopy

To compare EWI-2 surface expression between infected and uninfected cells, CEM-SS, CEM2n cells, and primary CD4^+^ T cells were infected with VSV-G-pseudotyped NL-sfGI as described above. Two to three days post infection, 3 × 10^5^ infected cells were plated onto each well of 8-well glass-bottom plates coated with 1:10 poly-L-Lysine in ddH_2_O. Two additional wells were used for uninfected controls. After 2 h of incubation at 37 °C, the medium was replaced with ice cold RPMI/10% FBS containing mouse anti-EWI-2 mAb at 1:200 dilution for surface labeling, and incubated for 45 min at 4 °C. Following the primary antibody incubation, cells were washed with RPMI/10% FBS and fixed with 4% PFA in PBS for 10 min at 4 °C, blocked and permeabilized with PBS containing 1% BSA and 100 μg/mL digitonin for 10 min, and incubated with the indicated secondary antibody in block for 45 min at room temperature. Cells were washed with block and imaged in PBS. At least 50 fields containing infected cells were selected for each biological replicate and imaged, deconvolved, and cropped using the DeltaVision microscope and Softworx software described above. After deconvolution, Fiji was used to manually select the cell surface at the midline of each cell and the mean intensity of EWI-2-associated signal was quantified and subsequently subtracted by the mean intensity of an area that did not contain cells. Cell-cell contact sites were excluded from the quantification. Background subtracted intensity values of all cells were normalized to the average surface associated intensity of the entire uninfected cell population, internal controls contained in the same wells as infected cells, contained within respective biological replicates. This normalization allowed for direct comparison of surface expression trends between biological replicates that accounts for potential variation in protein labeling efficiency between replicates. The virus-associated fluorescent reporter channel was used to segregate measurements into uninfected and infected. The data shown in Figure 3B are pooled from 2-3 independent biological replicates, each consisting of 2 technical replicates, all of which were sampled randomly until a minimum of 50 infected cells were quantified.

To compare EWI-2 surface expression levels between mononucleated infected cells and HIV-1-induced syncytia, primary CD4^+^ T cells were infected with VSV-G-pseudotyped virus as described above. Three days post infection, 3 × 10^5^ infected cells were plated onto each well of 8-well glass-bottom plates coated with 1:10 poly-L-Lysine in ddH_2_O alongside two wells of uninfected cells as controls. Cells were incubated at 37 °C for 2 h and surface labeled as described above using either mouse anti-EWI-2 or mouse anti-CD81 mAb at 1:200 or 1:100, respectively. Samples were fixed, permeabilized, and labeled with appropriate AlexaFluor conjugated antibodies and DAPI as described above. Cells were imaged in PBS and at least 50 fields containing 10-20 cells each, and containing at least some infected cells with multinucleated appearance (determined by DAPI and GFP signal) were selected for each biological replicate and imaged, deconvolved, and cropped as described above. Fiji was then used to analyze the surface expression of each protein of interest as described above. The virus-associated fluorescent reporter channel (GFP) was used to segregate measurements into infected and uninfected populations, and nuclear staining (DAPI) was used to further segregate infected cells into mononucleated and multinucleated infected cells. The EWI-2/CD81 channel was not viewed at all during imaging and field selection, or throughout image processing. The data shown in Figure 5 are pooled from 2-3 biological replicates, with two technical replicates each, all of which were sampled randomly until a minimum of 15 syncytia per biolobical replicate were quantified.

### 2.8 Determining surface EWI-2 signal on infected cells by flow cytometry

CEM2n cells infected as described above were harvested after three days and incubated in cold PBS with 5 mM EDTA for 15 min (3.0 × 10^5^ cells/tube). Cells were pelleted at 400 rcf for 7 min at 4 °C and resuspended in cold RMPI/10% FBS containing mouse anti-EWI-2 mAb at 1:200 dilution. After a 45 min incubation at 4 °C, cells were washed with cold RPMI/10% FBS and resuspended in ice cold PBS with 5 mM EDTA. To fix, an equal volume of PBS with 8% PFA was added and samples were incubated on ice for 10 min. Cells were washed and stained with Alexa Fluor 594-conjugated secondary antibody at 1:500 in block for 45 min at room temperature, before being washed, resuspended in PBS, and analyzed using a BD LSRII flow cytometer. Data were analyzed using FlowJo V10 (Becton, Dickinson & Company, Franklin Lakes, NJ). Samples were gated for infected and uninfected populations by GFP expression. EWI-2^high^ and EWI-2^low^ gates were set based in part on controls lacking primary antibody, and in part by adjusting the gates to reflect the number of uninfected EWI-2^high^ cells as measured by microscopy. The data shown are the collection of 3 independent biological replicates, each consisting of 2 technical replicates.

### 2.9 Establishment of EWI-2 knockdown CEM-SS cells

The shRNA-encoding sequenes targeting either EWI-2 (modified from previously described EWI-2-targeting siRNA [27] or a scrambled control, were introduced to the lentiviral vector FG12 (as described in 2.3) using oligos containing shRNA sequences, a loop sequence, and an AgeI site, flanked by BbsI and XhoI restriction site overhangs, as previously described [24], (EWI-2 sense, 5’-ACCGGGGCTTCGAAAACGGTGATCTTCAAGAGAGATCACCGTTTTCGAAGCC**C**TTTTTTACCGGTC-3’, and anti-sense, 5’-TCGAGACCGGTAAAAAAGGGCTTCGAAAACGGTGATCTCTCTTGAAGATCACCGTTTTCGAAGCCC-3’; scramble sense, 5’-ACCGGGCAGATGCGTCCAGTTAGATTCAAGAGATCTAACTGGACGCATCTGC**C**TTTTTTACCGGTC-3’, and anti-sense, 5’-TCGAGACCGGTAAAAAAGGCAGATGCGTCCAGTTAGATCTCTTGAATCTAACTGGACGCATCTGCC-3’). A PolII promoter was first obtained by ligating the oligo with PBS-hU6 digested with BbsI and XhoI rescrition endonucleases (New England BioLabs, Ipswich, MA). The PolII-shRNA constructs were obtained by digesting the resulting PBS-hU6 vector with XbaI and XhoI, and the insert was subsequently ligated into the FG12 vector digested with the same enzymes.

VSV-G pseudotyped FG12-shRNA lentiviruses were used to transduce CEM-SS cells by spinoculating one million cells with 500 μL of lentiviral supernatant (either shEWI-2 or shScramble). Cells were incubated at 37 °C for two-days in RPMI/10% FBS and positively transduced cells were then selected for puromycin resistance by supplementing the media with 0.5 μg/mL of puromycin for 8 days. shEWI-2 and shScramble CEM-SS cells were subsequently maintained in RPMI/10% FBS/0.25 μg/mL puromycin.

EWI-2 knockdown was analyzed by flow cytometry and microscopy. For flow cytometry analysis,

3.0 × 10^5^ shScramble and shEWI-2 cells, alongside parental CEM-SS controls, were pelleted at 400 rcf for 7 min, resuspended in 1:1000 Live/Dead Fixable Near-IR stain (ThermoFisher Scientific) in PBS for 30-45 min, washed with RPMI/10% FBS and fixed for 10 min in 4% PFA in PBS by resuspending the cells in PBS and then adding an equal volume of 8% PFA in PBS. Fixed samples were washed with 1 mL of PBS, blocked and permeabilized in 100 μL of block/perm buffer for 10 min, and washed with PBS containing 1% BSA. EWI-2 was labeled using mAb 8A12 diluted 1:200 in block for 45 min, washed with block, and stained with Alexa Fluor 488-conjugated secondary antibody in block for 45 min. Cells were then washed and resuspended in PBS for flow cytometry analysis using a BD LSRII flow cytometer. Data were analyzed using FlowJo V10. Samples were gated for live cells, and EWI-2 expression was measured by the mean fluorescence intensity of EWI-2 signal in the live cell population and normalized to the parental control expression within each biological replicate. Data are the result of 3 independent biological replicates with 2 technical replicates each. For microscopy, 2.5 × 10^5^ shScramble and shEWI-2 cells, alongside parental CEM-SS controls, were plated on 8-well glass bottom plates coated with 1:10 poly-L-lysine in ddH_2_O. After 2 h at 37 °C, cells were fixed for 10 min using 4% PFA in PBS, washed, and incubated with block/perm for 10 min. Cells were washed with block and incubated with 1:200 mAb 8A12 for 45 min, washed, and stained with 1:500 Alexa Fluor 647-conjugated secondary antibody and 1:2500 DAPI in block for 45 min. Cells were washed with block and imaged in PBS using a 60× objective as described above. Images were deconvolved and cropped by DeltaVision microscope and Softworx software described above and imported into Fiji for analysis.

### 2.10 CEM-luc-based HIV-1-induced cell-cell fusion assay

Two million shScramble or shEWI-2 cells were spinoculated as described above with 1.7 or 2 μL of VSV-G pseudotyped NL4-3, alongside parental CEM-SS cells spinoculated with 25 μL of VSV-G pseudotyped NL4-3 ΔEnv to achieve an infection rate of ~30% for each condition. Cells were incubated at 37 °C for 2 days and then co-cultured with uninfected CEM-luc cells in RPMI/10% FBS containing the following drug treatments; 1:1000 DMSO for vehicle control, 1 μM Efavirenz (EFV) (NIH AIDS Reagent Program, Cat. #4624) to inhibit transmission, or 1 μM EFV with 0.5 μM HIV-1 IIIB C34 peptide (C34) (NIH AIDS Reagent Program, Cat. #9824) to inhibit both transmission and cell-cell fusion. 24 h later, the co-culture medium was refreshed and all conditions were incubated at 37 °C in RPMI/10% FBS containing 1 µM EFV and 0.5 µM C34. 24 h later, cells were pelleted at 1000 rcf for 5 min at 4 °C and resuspended in luciferase reporter lysis buffer (Promega, Cat. #E4530) with 1% protease inhibitor cocktail (Millipore Sigma, Darmstadt, Germany, Cat. #P8340) to lyse on ice for 15 min. Lysates were cleared by centrifugation at 20,000 rcf for 5 min at 4 °C and stored at −80 °C until use for luciferase activity assays.

In parallel, infected cells were prepared for flow cytometry analysis alongside uninfected controls, to determine the infection rate across each condition at the start of the co culture with uninfected CEM-luc cells. Cells were pelleted and resuspended in 1:1000 Live/Dead Fixable Near-IR stain in PBS as described above, washed and resuspended in PBS. An equal volume of 8% PFA in PBS was added to fix the cells in a final concentration of 4% PFA in PBS for 10 min. Cells were washed and resuspended in block/perm, incubated for 10 min, washed with block, and resuspended for an overnight incubation in 1:100 AG3.0 in block. Cells were washed and stained with 1:500 Alexa Fluor 488-conjugated secondary antibody for 45 min followed by a wash with block. Cells were resuspended in PBS and analyzed by flow cytometry using a BD LSRII flow cytometer. Data was analyzed using FlowJo V10. Live cells were gated using the Live/Dead signal, and the percentage of infected cells in the live population was determined by gating on the AG3.0 associated signal.

Each lysate was incubated with an equal volume of firefly luciferase reagent (Promega, Cat. #E1500) for 1 min in a 96-well white-walled plate (ThermoFisher Scientific, Waltham, MA, Cat. #7571) before collecting luminescence signal intensity on a microplate reader (BioTek Synergy 2). Background luminescence was determined using a lysis buffer blank and subtracted from all experimental samples. Relative luminescence units (RLUs) were normalized based on the infection level of each cell type determined by flow cytometry analysis, and the average RLU value from the ΔEnv infected, DMSO treated condition was subtracted from all conditions. All samples treated with both EFV and C34 had RLU values below that of the ΔEnv DMSO condition (data not shown), validating the efficacy of the inhibitors for complete inhibition of transmission to target CEM-luc cells. To determine the proportion of luciferase expression due to cell-cell fusion, the average RLU value from the EFV-treated condition (syncytium formation-dependent signal) was divided by that of the DMSO-treated (signal from both transmission and syncytium formation) and multiplied by 100. Data represent the percentage of luciferase signal due to syncytium formation between infected shScramble or shEWI-2 cells and uninfected CEM-luc cells from 3 independent biological replicates each consisting of 1-2 technical replicates.

### 2.11 HeLa-based HIV-1-induced cell-cell fusion assay

50,000 HeLa cells were plated in each well of a 24-well plate and, the next day, transfected (using FuGENE6; see section 2.5) in duplicate with 100 ng of either pNL-sfGI or pNL-sfGI ΔEnv along with 500 ng total expression vector carrying CD81 or EWI-2. L6, a tetraspanin-like protein that does not inhibit cell-cell fusion, was co-transfected instead of CD81 or EWI-2 as a positive control for maximum fusion activity, For dose response assays, 125, 250, or 500 ng of either EWI-2 or CD81 plasmid was “stuffed” with L6 expression plasmid to maintain 500 ng of total protein expression plasmid in each condition. No cytotoxicity was observed upon transfection for any of the experimental conditions. 24 h post-transfection, producer HeLa cells were co-cultured with 10^6^ TZM-bl target cells (which, upon producer-target cell fusion, express firefly luciferase under control of the HIV-1 LTR) per well for 3 h before unattached target cells were washed off and the medium was refreshed. 14-18 h later, cells were lysed for at least 30 min on ice using 1% Triton X-100, 2mM EDTA, 50 mM Tris-HCl, 200 mM NaCl, with 1% protease inhibitor cocktail. Lysates were precleared by centrifugation at 20,000 rcf for 5 min at 4 °C and stored at −80 °C until use for luciferase activity assays. Note that the timepoints used here ensure that there is not enough time for the development of any luciferase signal resulting from productive infection of target TZM-bl cells through virus transmission (unpublished observation) and that only cell-cell fusion contributes to the luciferase activity measured.

Each lysate was incubated with an equal volume of firefly luciferase reagent for 1 min before collecting luminescence signal intensity on a microplate reader as described above (2.9). Background luminescence was determined using a lysis buffer blank and subtracted from all experimental samples. Luminescence intensity was used as a quantitave measurement of relative HeLa-TZM syncytium formation against the non-fusogenic (therefore incabable of forming syncytia) ΔEnv control by dividing each value by the ΔEnv value (which effectively corresponds to any leaky expression of luciferase in TZM-bl cells as no cell-cell fusion occurs at all in this condition). To then determine relative fusion activity of cells transfected with EWI-2 and CD81, those values were normalized to the L6 condition. Normalized fusion is therefore the fold difference of cell-cell fusion activity taking place when cells were co-transfected with the indicated amount of either CD81 or EWI-2 plasmid, compared to the activity taking place when cells were co-transfected with L6. The data shown are the collection of 4 independent biological replicates.

### 2.12 Statistical Analysis

All statistical analyses were carried out in GraphPad Prism 8 as indicated in Figure legends.

## 3. Results

### 3.1 EWI-2 accumulates at the virological presynapse in HIV-1-infected cells

Because EWI-2 is known to associate with ezrin and CD81 [25,27], two cellular factors that accumulate at the producer cell side of the virological synapse (VS) [24,54], we first sought to determine whether this protein also localizes to the VS. CEM-SS cells were infected with (VSV-G-pseudotyped) NL4-3 WT or NL4-3 ΔEnv (virus that does not express Env) and mixed with target CEM-SS cells (labeled with a cytoplasmic dye). Upon imaging with a 60× objective, the VS was identified and defined by region selection as clusters of immunolabeled Gag present at producer-target cell contact sites. DIC was used to identify and region-select cell-cell contacts between ΔEnv producers and uninfected target cells as Gag will not accumulate at these contacts in the absence of Env [1]. The EWI-2 channel was not viewed during the process of defining VS/contact regions to eliminate possible bias. To calculate enrichment at the VS/contact, we divided the EWI-2 signal intensity within the defined VS/contact site by the sum of the EWI-2 surface intensity at non-contact sites on the producer and target cell at each VS/contact. This unbiased approach prevents potential inflation of the enrichment value that could occur if we assumed that EWI-2 was solely contributed by either the target or producer cell. Similarly to p-ezrin and CD81 [24,54], EWI-2 was observed to co-accumulate with Gag at the VS in an Env-dependent manner (Figure 1A). EWI-2 signal intensity was ~4-fold enriched at the VS in CEM-SS cells infected with NL4-3 WT, while no EWI-2 enrichment was seen at cell-cell contacts in cells expressing NL4-3 ΔEnv (Figure 1A). EWI-2 signal intensity was also enriched ~1. 6-fold at the VS in infected primary CD4^+^ T cells at Env-dependent VSs, and was again not enriched at non-VS contact sites (ΔEnv) (Figure 1B).

**Figure 1.**
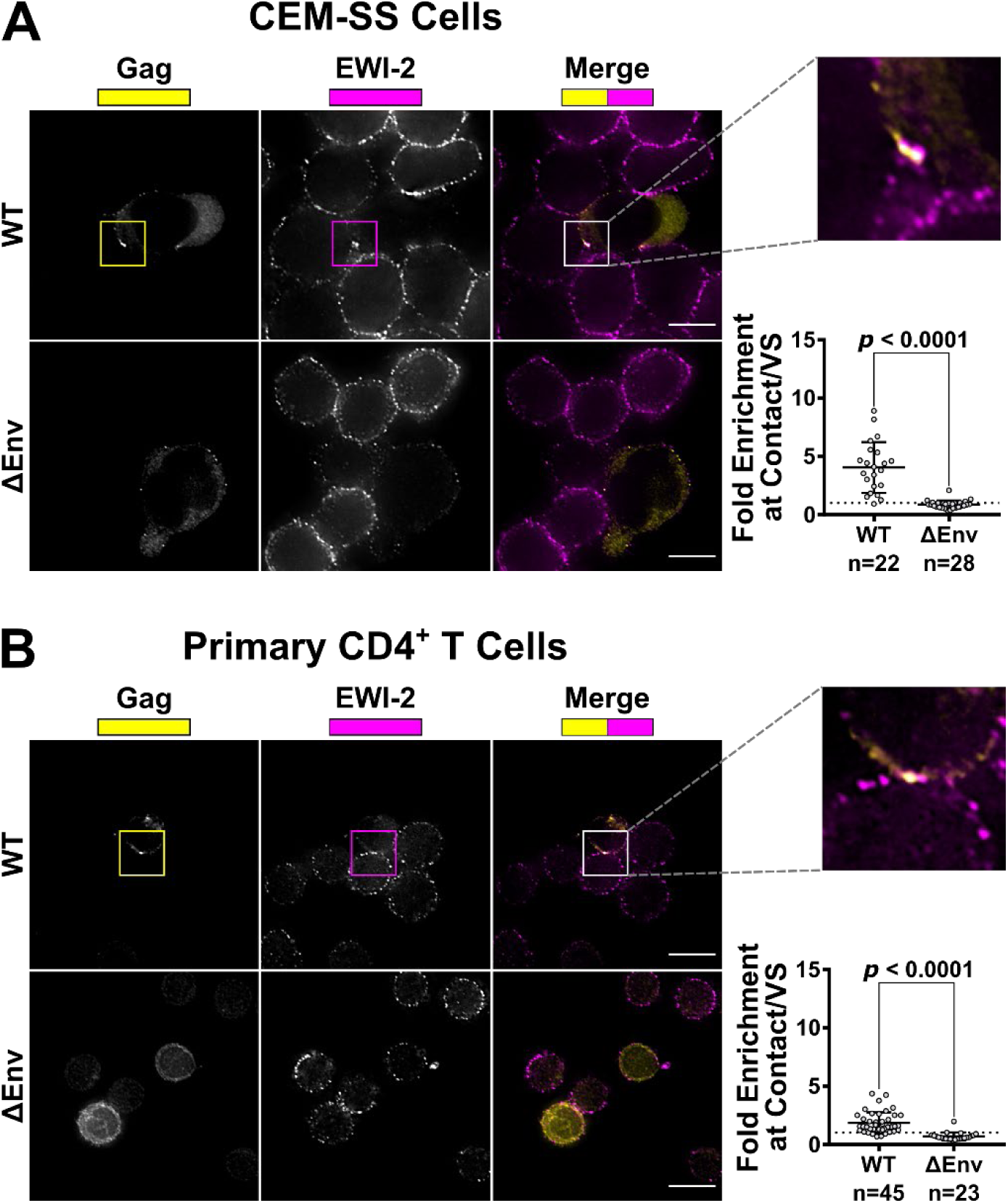
EWI-2 co-accumulates with Gag at the HIV-1 VS in T cells. (**a**) CEM-SS cells infected with HIV-1 NL4-3 WT or ΔEnv were co cultured with uninfected CEM-SS target cells for 5 h, and subsequently stained for surface EWI-2 (magenta) and Gag (yellow). The EWI-2-associated fluorescence intensity at cell-cell contacts either enriched with Gag (WT) or not Gag-enriched but identified by DIC (ΔEnv) was measured. This value was then divided by the sum of the EWI-2-associated fluorescence intensity on non-contact sites on the producer and target cell in each VS/contact to yield EWI-2 enrichment (i.e. the values shown here). The data quantified are from one biological replicate consisting of two technical replicates. Similar trends were observed in a second dataset; not shown. (**b**) Primary CD4^+^ T cells infected with NL-sfGI WT or NL-CI ΔEnv were co-cultured with uninfected target primary cells for 2 h and stained for EWI-2 (magenta) and Gag (yellow), followed by secondary pAbs (Alexa Fluor 647-conjugated for EWI-2, and either Alexa Fluor 594 or Alexa Fluor 488-conjugated for Gag in the case of WT and ΔEnv, respectively). Because different secondary antibodies were used for Gag in either condition, the scaling shown for that channel is not the same across the two conditions, and was based on corresponding no primary and uninfected controls done alongside each dataset. Enrichment of EWI-2 at Env-dependent (WT) or Env-independent (ΔEnv) infected-uninfected cell contacts was quantified as described in (**a**). The data quantified are pooled from two independent biological replicates, each consisting of two technical replicates. Scale bars = 10 μm. In both data plots, each data point represents one cell-cell contact site (as opposed to one cell). The dotted horizontal line indicates a theoretical fold enrichment value of 1, which indicates no enrichment. Error bars = standard deviation of the mean (SD). *p*-values are the result of two-tailed non-parametric Mann-Whitney *U* tests.

To determine whether EWI-2 enrichment at the VS takes place within the infected cell, i.e. at the presynaptic terminal (rather than the apposed uninfected target cell), HIV-1-infected CEM-SS cells were co-cultured with uninfected target TZM-bl cells (which have nearly-undetectable levels of EWI-2 on their surface) and imaged as described above. Significant EWI-2 enrichment (~5.3-fold) was observed at the VS as before (Figure 2A), demonstrating that the observed EWI-2 accumulation in CEM-SS-CEM-SS co-cultures takes place at least partially within the producer cell. To evaluate the relative contribution of any postsynaptic (i.e. target cell-side) accumulation of EWI-2, HIV-1-producing HeLa cells (which, like TZM-bl cells, also exhibit nearly-undetectable levels of EWI-2 on their surface) were cocultured with uninfected target CEM-SS cells. In this case, minimal EWI-2 accumulation was detected at synapses (~1.1-fold; Figure 2B), showing that EWI-2 enrichment seen at T cell-T cell VSs takes place (almost) exclusively at the presynaptic terminal of the VS, i.e. in the producer cell. Together, these results conclusively document that EWI-2 is recruited to the virological presynapse during HIV-1 cell-to-cell transmission.

**Figure 2.**
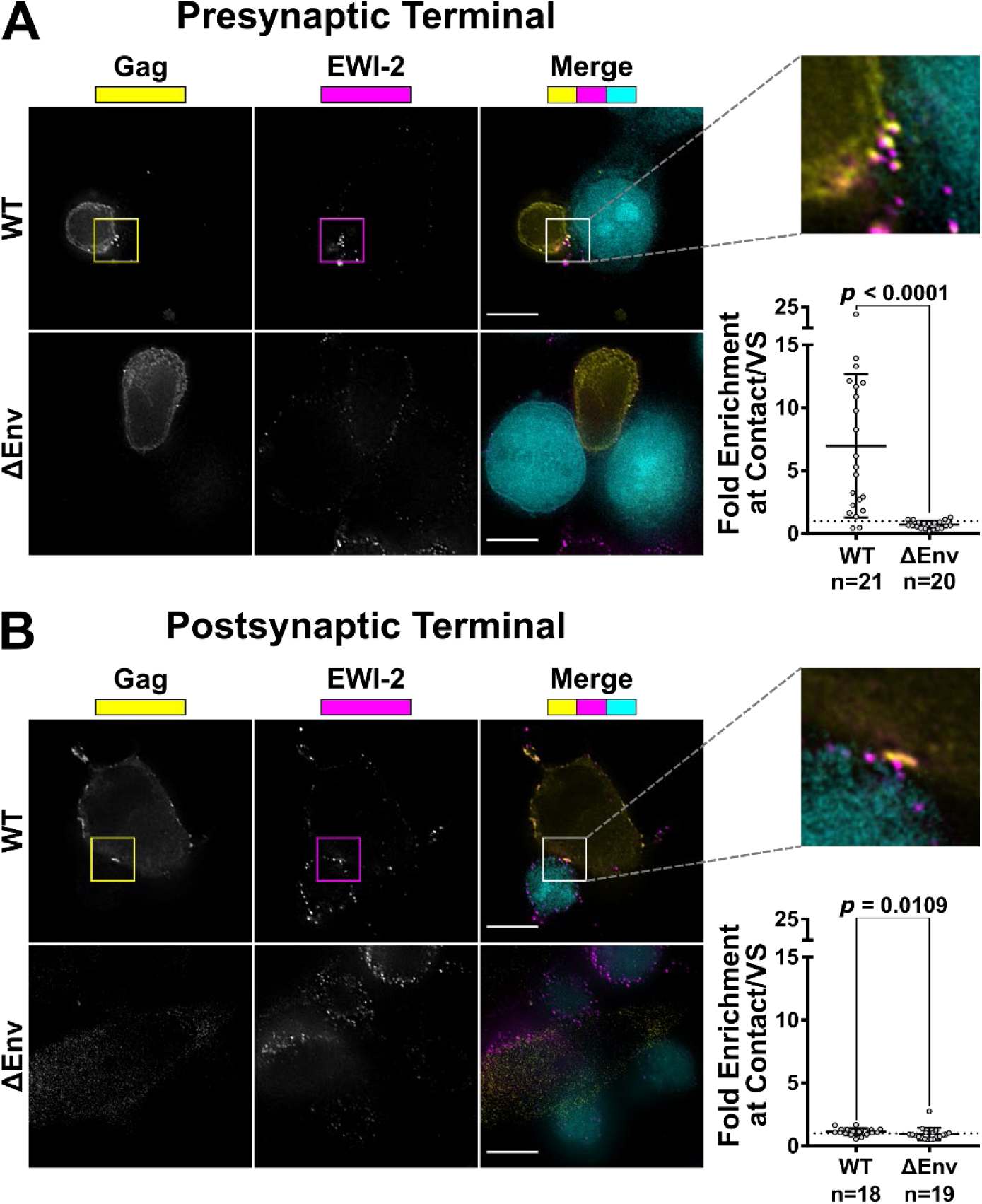
EWI-2 accumulation takes place on the producer cell side of the VS. (**a**) To evaluate presynaptic accumulation of EWI-2, CEM-SS cells infected with HIV-1 NL-CI WT or ΔEnv were co cultured with CMAC (cyan) labeled TZM-bl target cells (which have nearly-undetectable EWI-2 surface levels compared to CEM-SS cells) for 2.5 h, and subsequently stained for surface EWI-2 (magenta) and Gag (yellow). EWI-2 enrichment was quantified as described in Figure 1. Quantification is the result of pooled VS/contacts from two independent biological replicates. (**b**) To evaluate postsynaptic accumulation of EWI-2, HeLa cells (which, like TZM-bl cells, also have nearly-undetectable EWI-2 surface levels) were transfected with HIV-1 NL-sfGI or NL-sfGI ΔEnv and cocultured with uninfected CEM-SS target cells (cyan) for 2-2.5 h. Cells were stained for surface EWI-2 (magenta) and Gag (yellow). Note that Gag expression in the ΔEnv condition was quite low, since Gag expression in this virus is already expected to be considerably reduced [55]. EWI-2 enrichment was calculated as described in Figure 1. Quantification is the result of pooled VSs/contacts from two independent biological replicates. Scale bars = 10 μm. In both data plots, each dot represents the EWI-2 enrichment value of one VS/contact. The dotted horizontal line indicates a theoretical fold enrichment of 1, which indicates no enrichment. Error bars = standard deviation of the mean (SD). *p*-values are the result of two-tailed non-parametric Mann-Whitney *U* tests.

### 3.2 Overall surface levels of EWI-2 are decreased upon HIV-1 infection

Despite its enrichment at the virological presynapse, the EWI-2 partner protein CD81 (as well as other tetraspanins) is overall downregulated in HIV-1-infected cells [54,56,57]. We previously used Tandem Mass Tag (TMT)-based quantitative proteomics to map global changes in whole cell protein abundances in HIV-infected T cells [50,51]. Like CD81, EWI-2 was decreased in abundance in both CEM-T4 T cells and primary human CD4^+^ T cells (Figure 3A). To confirm these data using an orthogonal approach, we tested whether surface levels of EWI-2 are decreased in lymphocytes infected with HIV-1 NL-sfGI, a strain in which superfolder GFP (sfGFP) replaces the Nef gene and Nef expression is restored using an IRES [10]. We chose to utilize this GFP reporter virus, rather than immunolabeling Gag after fixation, because Gag-negative (or undetectable) cells still in the early phase of infection may exhibit host protein downregulation due to early Nef expression (reviewed in [58]).

**Figure 3.**
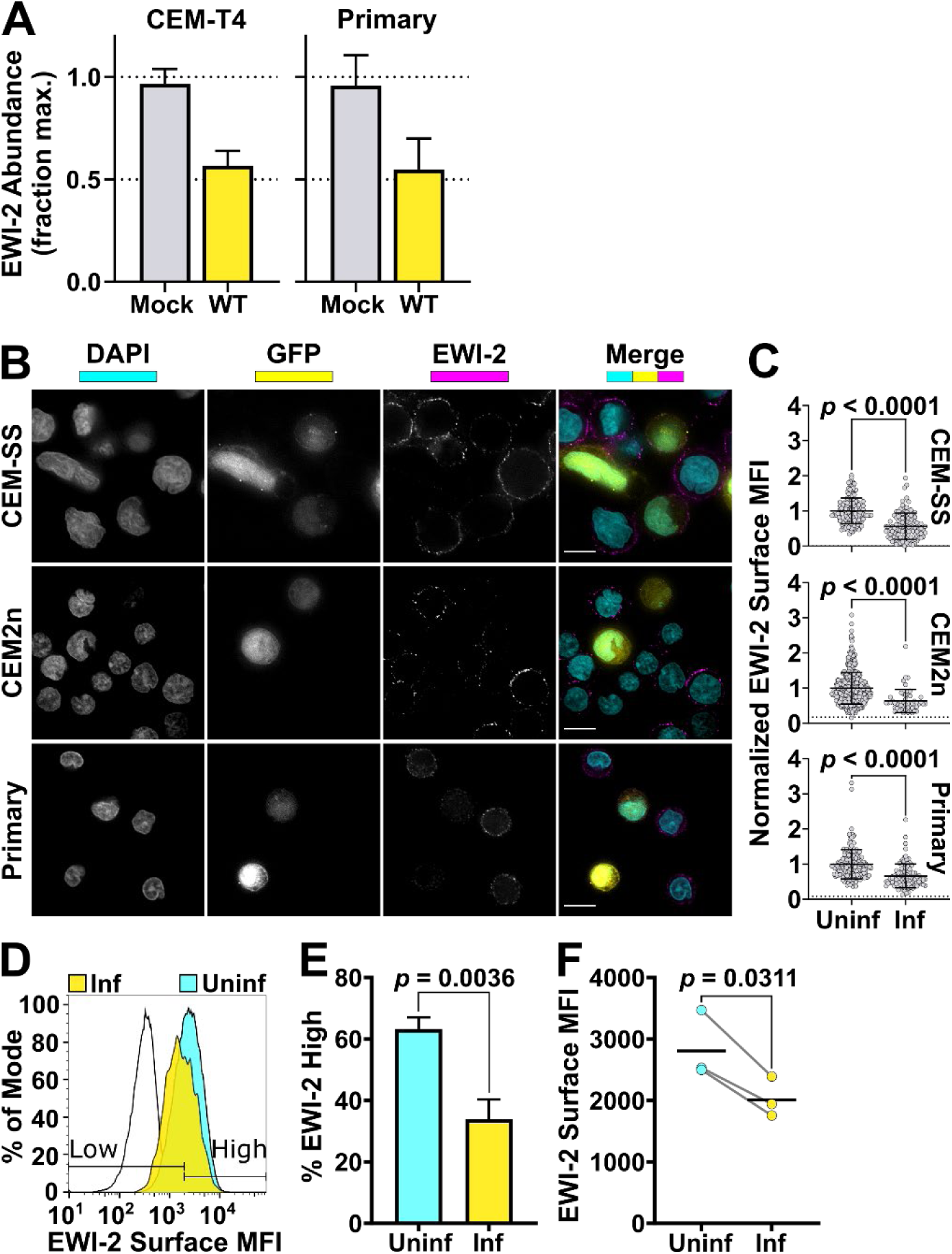
EWI-2 is downregulated from the surface of infected cells. (**a**) Abundance of EWI-2 in mock-infected (gray) versus WT HIV-infected (yellow) CEM-T4 T cells or primary human CD4^+^ T cells. Experiments were conducted in triplicate and whole cell lysates subjected to Tandem Mass Tag (TMT)-based quantitative proteomics 48 h after infection (reanalysis of data from [50] and [51]). 7 (CEM-T4 T cells) or 6 (primary human CD4+ T cells) unique peptides were used for EWI-2 quantitation. Mean relative abundances (fraction of maximum TMT reporter ion intensity) with 95% confidence intervals (CIs) shown. (**b**) Cells were infected with NL-sfGI and surface-labeled for EWI-2, fixed, stained with DAPI (shown in cyan) and Alexa Fluor 594-conjugated secondary antibody, and imaged. GFP signal (yellow) was used to identify infected cells, and EWI-2-associated signal is shown pseudocolored in magenta. Representative cells are shown. Scale bars = 10 μm. (**c**) Cells were prepared as in (**b**) and EWI-2 levels at the plasma membrane in infected (Inf) and uninfected (Uninf) cells were measured by manually selecting the plasma membrane at the midline of each cell and quantifying the mean EWI-2-associated fluorescence intensity. Fluorescence intensity of each cell was normalized to the average intensity value of uninfected cells within the same imaging set. Data shown are pooled from two to three biological replicates, each consisting of two technical replicates. Only non-contact sites were quantified. Error bars = SD. *p*-values are the result of a two-tailed non-parametric Mann-Whitney *U* test. (C-E) CEM2n cells were infected with NL-sfGI and surface-labeled for EWI-2, fixed, and stained with Alexa Fluor 647-conjugated secondary antibody, and analyzed by flow cytometry. (**d**) Representative histogram normalized to mode of the EWI-2 signal intensity at the cell surface for unstained controls (black outline), infected cells (yellow), and uninfected cells (cyan). The gates defining EWI-2^high^ and EWI-2^low^ cells are shown. (**e**) Data represent the percentage of uninfected and infected cells that fell into the EWI-2^high^ gate shown in (**d**) from 3 independent biological replicates, averaged across 2 technical replicates within each. (**f**) EWI-2 surface expression was measured by mean fluorescence intensity (MFI) of EWI-2-associated signal. In both panels, lines connect paired data points, i.e. infected cells and uninfected cells (within an infected tube) from the same biological replicate. Error bars = SD. *p*-values (E-F) are the result of a two-tailed paired *t* test.

HIV-1-infected cells adhered to glass-bottom dishes were surface-labeled with EWI-2 primary antibody on ice, and fixed before incubation with fluorescent secondary antibody. Uninfected and HIV-1-infected cells were imaged with a 60× objective and the resulting images were deconvolved. The mean fluorescence intensity (MFI) of EWI-2 on the surface of each cell was determined by measuring the EWI-2-associated signal intensity of manually-selected regions of the cell surface (representative images shown in Figure 3B) and normalizing the raw MFI of each cell to the average EWI-2 signal from uninfected cells within the same imaging set. After measuring surface MFI, on average across three independent biological replicates, infected (GFP-expressing) cells had significantly lower (~2-fold) EWI-2-associated signal than uninfected (GFP-negative) cells, after subtracting background signal (Figure 3B). This phenomenon was consistent across CEM-SS, CEM2n, and primary CD4^+^ T cells.

We also sought to quantify EWI-2 surface expression by flow cytometry as a means of high-throughput analysis. HIV-1 NL-sfGI-infected CEM2n cells, surface-labeled for EWI-2 and analyzed by flow cytometry, were gated for high or low levels of EWI-2 using appropriate controls (representative histogram plots shown in Figure 3D). These data showed that a much lower proportion of infected cells (identified as GFP^+^) had high levels of EWI-2 surface expression than of uninfected cells (identified as GFP^−^) in the same culture (Figure 3E). Additionally, the mean fluorescence intensity of EWI-2-associated signal was lower within the total population of infected cells compared to that of the uninfected cells (Figure 3F).

Like other cell surface proteins downregulated by HIV-1, depletion of CD81 (as well as other tetraspanins) is mediated by the accessory proteins Vpu (predominantly) and Nef [56,57]. We have previously shown that substrates of different HIV-1 accessory proteins may be distinguished by their characteristic patterns of temporal regulation in HIV-1-infected T cells [50,51,53]. Accordingly, the temporal expression profile of plasma membrane EWI-2 was strikingly similar to that of BST2 (Tetherin), a canonical Vpu target (Figure 4A).

**Figure 4.**
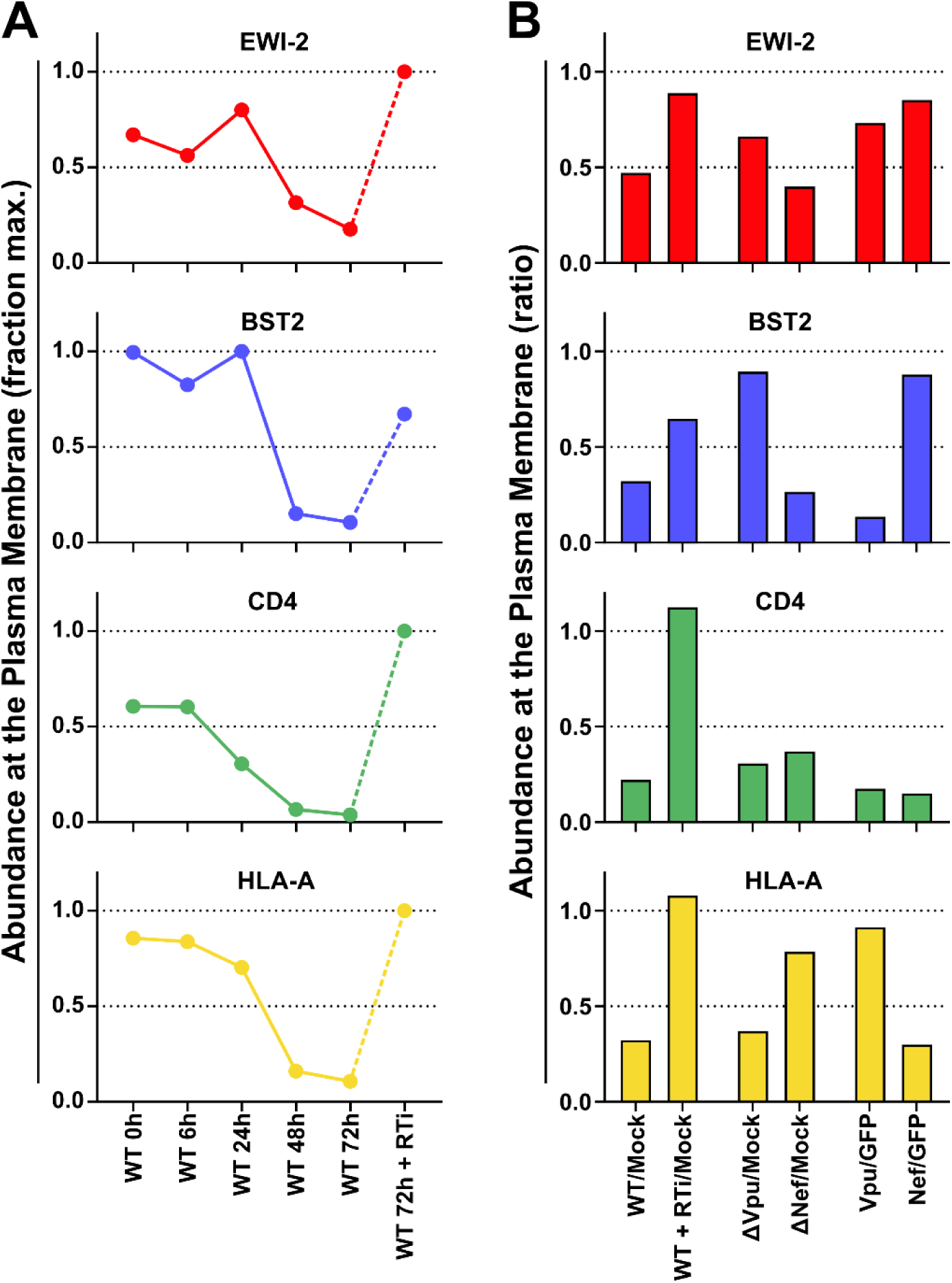
Plasma membrane EWI-2 is downregulated by Vpu. (**a**) Temporal expression profiles of cell surface EWI-2 (red, upper panel) or indicated control proteins (blue/green/gold, lower panels) in WT HIV-1-infected CEM-T4 T cells (reanalysis of data from [53]). Plasma membrane proteins were subjected to TMT-based quantitative proteomics 0 (uninfected), 6, 24, 48, and 72 h after infection, or 72 h after infection in the presence of reverse transcriptase inhibitors (RTi). 12 unique peptides were used for EWI-2 quantitation. Relative abundances (fraction of maximum TMT reporter ion intensity) are shown. (**b**) Abundance of EWI-2 (red, upper panel) or indicated control proteins (blue/green/gold, lower panels) in control CEM-T4 T cells or CEM-T4 T cells infected with WT HIV-1 in the presence/absence of RTi, infected with Vpu- or Nef-deficient HIV-1, or transduced with Vpu or Nef as single genes (reanalysis of data from [53]). Plasma membrane proteins were subjected to Stable Isotope Labelling with Amino acids in Cell culture (SILAC)-based quantitative proteomics 72 h after infection (3 x 3-way comparisons). 12 (WT HIV-1 +/− RTi), 9 (∆Vpu/∆Nef HIV-1) or 14 (Vpu/Nef) unique peptides were used for EWI-2 quantitation. Ratios of abundances to mock-infected CEM-T4 T cells (WT HIV-1 +/− RTi and ∆Vpu/∆Nef HIV-1) or GFP-transduced CEM-T4 T cells (Vpu/Nef) are shown.

Furthermore, like BST2, depletion of cell surface EWI-2 by HIV-1 infection was abrogated in the presence of reverse transcriptase inhibitors, and when cells were infected with Vpu-deficient HIV-1 (Figure 4B). Taken together, our proteomic data therefore strongly suggest that Vpu is primarily responsible for HIV-1-dependent EWI-2 downregulation. As with the tetraspanins, however, the incomplete rescue in the presence of Vpu-deficient virus, and relatively modest depletion when Vpu was expressed as a single gene (Figure 4B), suggest that Nef may also contribute to depletion of cell surface EWI-2 in the context of HIV-1 infection.

### 3.3 EWI-2 inhibits HIV-1-induced syncytium formation

Likely through their accumulation at the producer cell side of the VS, the EWI-2 partner proteins CD81 and ezrin repress fusion of infected and uninfected cells, i.e. syncytium formation [22–24]. Given that EWI-2 also accumulates at the VS (Figure 1), we sought to test whether it also contributes to the inhibition of HIV-1-induced syncytium formation by reducing its expression using RNA interference.

We established an EWI-2 knockdown CEM-SS cell line by lentiviral transduction using a targeting vector (FG12) that directs expression of a short hairpin RNA (shRNA) targeting EWI-2 (shEWI-2), using the same targeting sequence as in a previous report [32]. As a control, this targeting sequence was scrambled several times, all resulting sequences were tested against the human genome by BLASTn, and the sequence with the least homology to any human transcript was selected (shScramble, or shScr). This modified FG12 vector also carries a puromycin resistance cassette, while the GFP reporter cassette (as used in [24]) was removed to allow use of GFP reporter viruses. The puromycin-resistant shEWI-2 CEM-SS cells were analyzed by microscopy (Figure 5A) and by flow cytometry (Figure 5B-C), and were found to have ~3-fold reduced EWI-2 surface levels, compared to both the shScramble control and the parental non-transduced CEM-SS cells.

**Figure 5.**
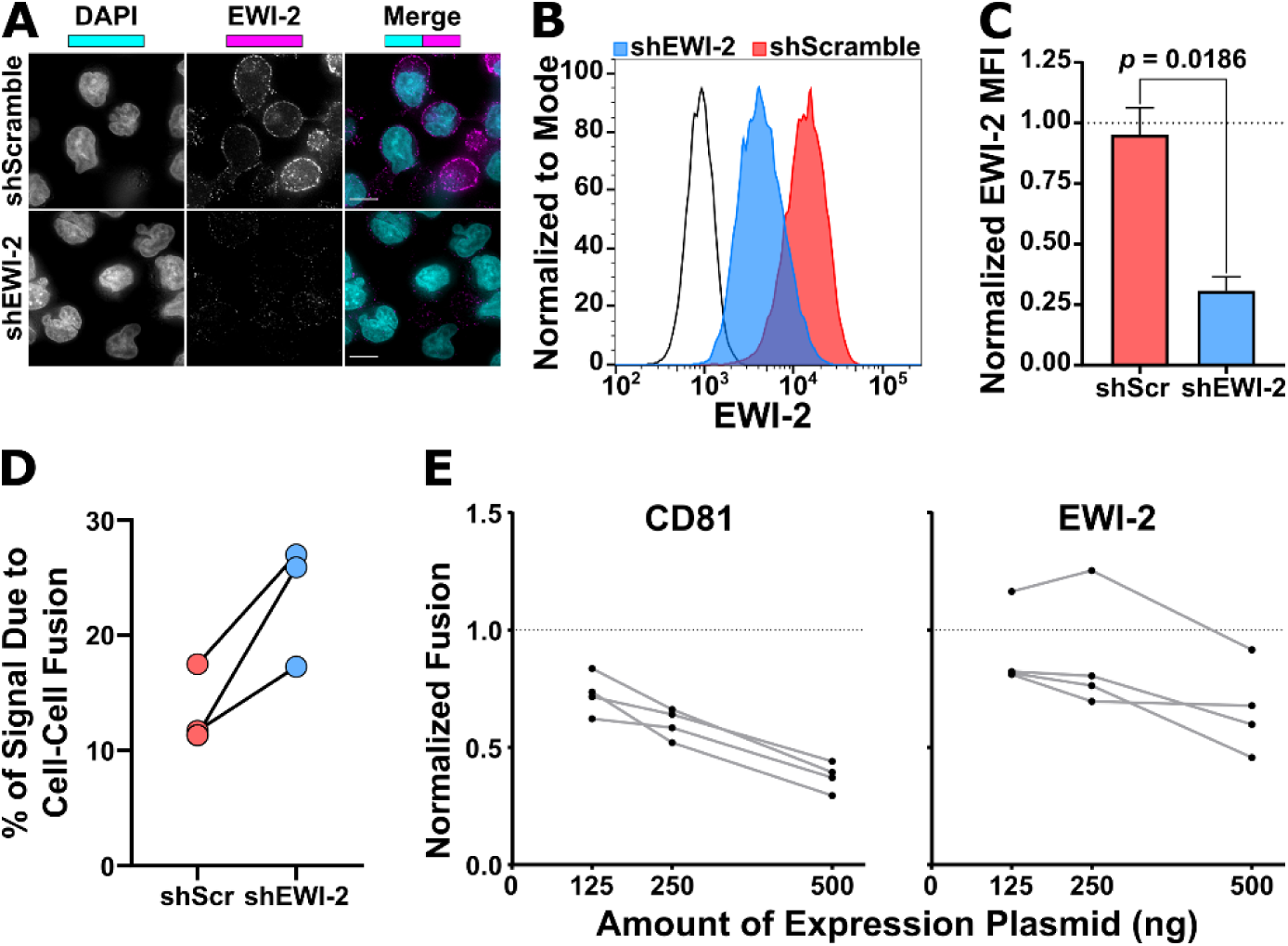
EWI-2 inhibits infected-uninfected cell fusion. (**a-c**) EWI-2 expression in shScramble (shScr) and shEWI-2 CEM-SS cells was analyzed by microscopy (**a**) and flow cytometry (**b-c**). (**a**) For microscopy, cells were plated onto poly-L-lysine-coated glass, fixed, permeabilized, labeled for EWI-2, and stained using fluorescent secondary antibody (magenta) and DAPI (cyan). (**b-c**) For flow cytometry analysis, cells were labeled with Live/Dead Fixable Near-IR, fixed, permeabilized, labeled for EWI-2, and stained with fluorescent secondary antibody. (**b**) Representative histogram of the EWI-2 signal intensity normalized to mode in live cells for unstained controls (black line), shEWI-2 (blue), and shScr (red) cells. (**c**) Average EWI-2 MFI in live shScr (red) and shEWI-2 (blue) cells from 3 independent biological replicates, normalized to EWI-2-labeled parental CEM-SS cells (represented at a value of 1 with a dashed line). Error bars = SD. *p*-value is the result of a paired *t* test. (**d**) CEM-luc fusion assays were performed using shScr or shEWI-2 producer cells infected with NL4-3, which were co-cultured with CEM-luc target cells in the presence of DMSO (vehicle control), EFV (luciferase signal resulting exclusively from cell-cell fusion), or EFV + C34 (to inhibit all transmission and cell-cell fusion) alongside parental CEM-SS cells infected with NL4-3 ΔEnv co-cultured with CEM-luc cells in the presence of DMSO. Luminescence readings (across 3 independent biological replicates) from the EFV-treated condition was divided by the DMSO reading from the same producer cell type and multiplied by 100 to determine the percentage of luciferase expression dependent on cell-cell fusion (syncytium formation) between either shScr or shEWI-2 producer and CEM-luc target cells. Values from the same biological replicate are linked by a black line. (**e**) HeLa-TZM-bl fusion assays were performed using producer HeLa cells that were co-transfected with either pNL-sfGI ΔEnv (ΔEnv) or pNL-sfGI (WT) in combination with either EWI-2, CD81, or L6 overexpression plasmids. Luminescence readings (across 4 independent biological replicates, each with 2 technical replicates) were divided by the ΔEnv condition to obtain the fold increase in fusion, and then normalized to the WT co-transfected with L6 condition (thus making L6 have a value of 1, shown as a dashed line). Values from the same biological replicate are linked by a grey line.

shEWI-2 and shScramble cells were then assayed for their ability to support HIV-1-induced cell-cell fusion with CEM-luc cells as target cells, using a previously reported assay that discriminates between the luciferase signal derived from active virus transmission and signal from cell-cell fusion [24,59]. Across three independent biological replicates, HIV-1-infected shEWI-2 cells were found to form syncytia considerably more frequently (~1.8-fold) than HIV-1-infected shScramble cells (Figure 5D).

In parallel, and as we have done previously to examine the fusion-inhibitory capacity of tetraspanins [22,23], we tested whether EWI-2 inhibits HIV-1-induced syncytium formation in a dose-dependent manner by overexpressing EWI-2 in HeLa cells (which have nearly-undetectable endogenous levels of EWI-2). NL-sfGI-producing HeLa cells overexpressing either EWI-2, CD81, or L6 (a tetraspanin-like surface protein that does not repress HIV-1-induced cell-cell fusion; [23,60]) were co-cultured with uninfected target TZM-bl cells. As a negative control for HIV-1-induced cell-cell fusion, Env-deleted (ΔEnv) NL-sfGI-expressing HeLa cells were also co-cultured with target TZM-bl cells. HIV-1-induced HeLa-TZM-bl syncytia express firefly luciferase under control of the HIV-1 LTR [22]. After 3 h of co-culture (and another 14-18 h to allow for reporter expression), cells were lysed, the lysates were incubated with luciferase substrate, and luminescence was measured using a microplate reader. Overexpression of increasing amounts of EWI-2 (125, 250, or 500 ng of plasmid) in NL-sfGI-producing cells resulted in robust and dose-dependent decrease of cell-cell fusion (at 250 and 500 ng of input plasmid), though repression was not as extensive as that observed upon CD81 overexpression (Figure 5E).

Taken together, the accumulation of EWI-2 at the presynaptic terminal of the HIV-1 VS (Figures 1–2), the concomitant overall downregulation of EWI-2 in infected T cells (Figure 3), and the requirement for high EWI-2 expression for efficient control of Env-induced cell-cell fusion (Figure 5) establish EWI-2 as a host fusion-inhibitory protein harnessed by HIV-1 during cell-to-cell virus transmission.

### 3.4 EWI-2 and CD81 surface expression is restored on HIV-1-induced syncytia

HIV-1-infected cells have been well documented to have altered surface expression profiles compared to uninfected cells (reviewed in [61]). However, previous analyses (including ours) were performed using bulk populations of HIV-1 infected cells, and thus could not or did not discriminate between mono- and multinucleated HIV-1-infected cells. HIV-1-induced syncytia likely have altered surface expression compared to mononucleated infected cells, as the process of syncytium formation (infected-uninfected cell fusion) provides a sudden influx of yet-to-be downregulated host proteins contributed by the uninfected target cell upon membrane merger and cytoplasm mixing. Therefore, we chose to use microscopy to analyze the surface expression of EWI-2 and CD81 on HIV-1-infected cells in order to, for the first time, confidently discriminate between mononucleated infected cells and multinucleated HIV-1-induced syncytia.

HIV-1-infected primary CD4^+^ T cells were cultured for three days post infection to allow time for syncytium formation. Infected cells were plated, surface-labeled for EWI-2 or CD81 on ice, and fixed prior to incubation with secondary antibody and imaging as before. The surface expression of each cell was quantified, normalized to internal uninfected controls, and data were segregated into populations of uninfected cells, mononucleated infected cells, and multinucleated infected cells (syncytia, identified as multinucleated by DAPI nuclear staining and positive for the viral reporter (GFP), as shown in representative images; Figure 6A). Strikingly, we found that syncytia had restored surface expression of both EWI-2 and CD81, back to nearly the same level as uninfected T cells found within the same wells (Figure 6B).

**Figure 6.**
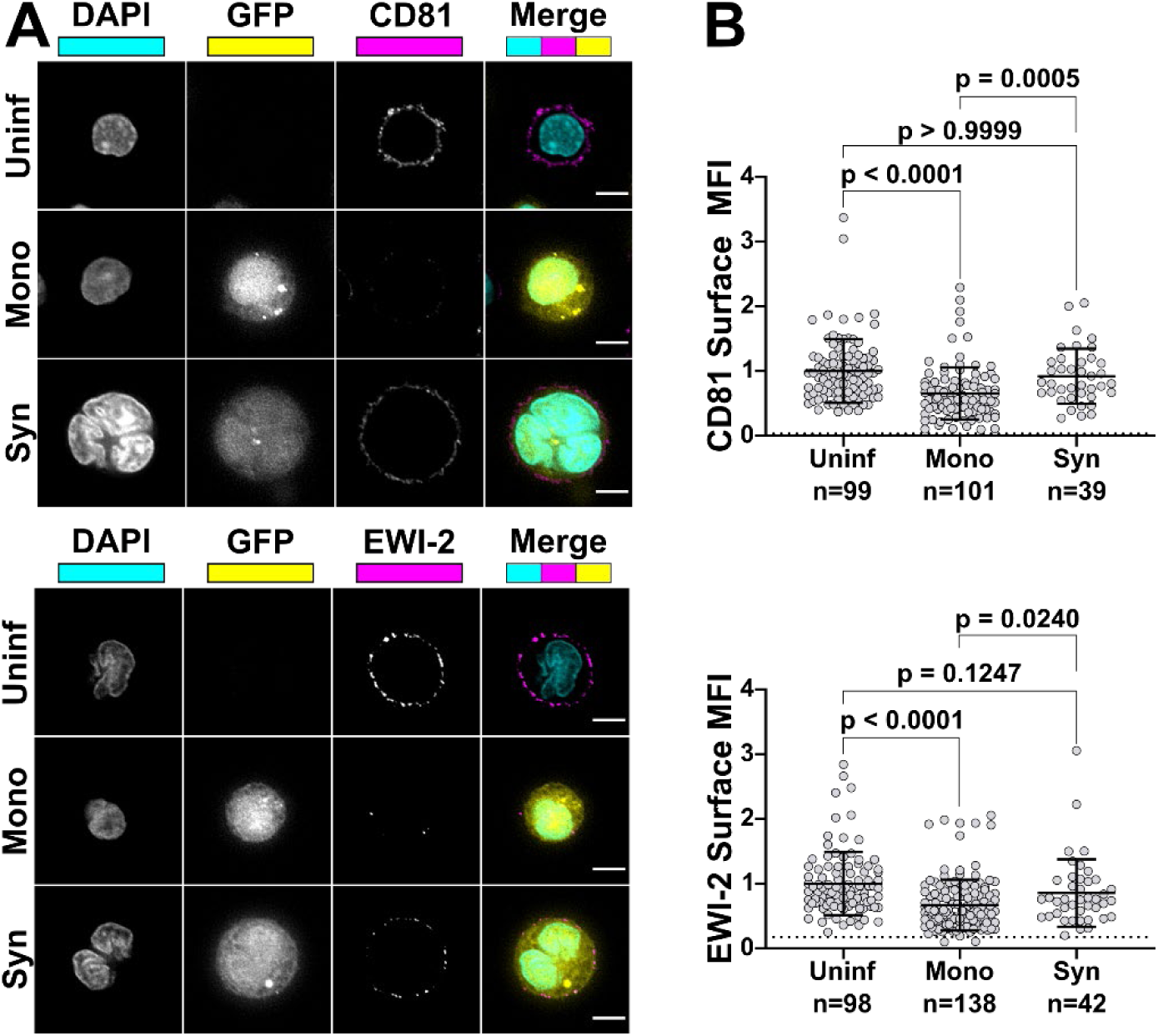
Syncytia have higher surface expression of EWI-2 and CD81 than mononucleated infected cells. (**a**) Primary CD4^+^ T cells were infected with NL-sfGI, surface-labeled for either EWI-2 or CD81 (both shown in magenta), fixed, stained with DAPI (cyan) and AlexaFluor 647-conjugated secondary antibody, and imaged. Infected cells were identified by GFP (yellow), and discriminated as mono- or multinucleated infected cells by DAPI. Representative cells are shown. Scale bars = 5 μm. (**b**) Cells were prepared as described in (**a**) and analyzed for EWI-2 or CD81 surface expression on uninfected cells, mononucleated infected cells (Mono) and syncytia (Syn) by manually selecting the plasma membrane at the midline of each cell and quantifying the mean EWI-2 or CD81-associated fluorescence intensity. Raw fluorescence intensity values were background-subtracted using the fluorescence intensity of a cell-free area within the same image and subsequently normalized to the average intensity value of uninfected cells within the same imaging set. Data shown are the pooled normalized intensity values of two independent biological replicates, each with two technical replicates. Each data point represents the normalized surface MFI of an individual cell. Error bars = SD. *p*-values are the result of two-tailed non-parametric Mann-Whitney *U* tests.

### 4. Discussion

Transient alignment of infected (producer) and uninfected (target) cells allows for efficient transmission of virus particles. However, because of the presence of viral Env and CD4/co-receptor at the surface of producer and target cell, respectively, rather than separating after particle transfer, these cells could also easily fuse with each other, thus forming a syncytium. This study now identifies EWI-2 as a host protein that contributes to maintenance of viral homeostasis through fusion inhibition.

Our investigations were partially prompted by two recent reports. In one of those studies, Rubinstein and colleagues documented a role for EWI-F, a close relative of EWI-2, in myoblast fusion regulation [26]. EWI-F was shown to act as fusion repressor in cooperation with the tetraspanins CD9 and CD81. With the other study, Yáñez-Mó and colleagues [32] showed the presence of EWI-2 at sites of contact between uninfected T cells and T cells stably expressing HIV-1 Env. In separate experiments, HIV-1-infected EWI-2 knockdown cells were also shown to have somewhat increased virus production and the authors mentioned (as data not shown) that this was accompanied by augmented syncytium formation, indicating that EWI-2 could be involved in the regulation of HIV-1-induced membrane fusion. Importantly, however, the study did not address the question of whether the reported increase in syncytium formation was (potentially) caused by the action of EWI-2 in producer or target cells, nor did it provide a dissection of where EWI-2 accumulates (producer and/or target cells). The authors did speculate that EWI-2, together with α-actinin, might be active in target cells, there possibly contributing to α-actinin’s actin bundling activity, thus ultimately inhibiting virus entry/fusion. They also explicitly stated, however, that even if their speculation about where α-actinin acts during virus replication should eventually turn out to be confirmed (with subsequent studies), they cannot exclude an involvement of the partner protein EWI-2 in “subsequent steps of the viral life cycle”. Our study now reveals that EWI-2 indeed acts during the late phase of the HIV-1 replication cycle: It accumulates on the producer cell side of the VS (Figures 1–2). Surprisingly, unlike tetraspanins, which have fusion-inhibitory roles at both sides of the VS (and thus are present at both the viral pre- and postsynapse [22,62]), EWI-2 accumulates (and inhibits fusion) only at the presynaptic terminal of the VS. This leads us to speculate whether EWI-2 accumulation at the presynaptic terminal might contribute to unique intracellular signaling events in HIV-1-infected cells [32,63], such as tuning T cell receptor function.

Paralleling what we previously documented for tetraspanins [22], we found that fusion with uninfected target cells was inhibited by EWI-2, and we established that it does so in a dose-dependent manner (Figure 5). Also analogous to our findings about tetraspanins [54,56], we demonstrate that while EWI-2 accumulates at the virological presynapse, overall this protein is downregulated in infected cells (Figure 3). Our proteomic analysis (Figure 4) now shows that EWI-2 depletion from the infected cell surface, as is also the case for tetraspanins [56,57], is primarily mediated by Vpu (Figure 4). Since EWI-2 is a known interactor of tetraspanins CD81 and CD9, it is possible that EWI-2 downregulation by Vpu (with or without Nef) is “direct” (like e.g. the canonical Vpu “targets” BST2 and CD4, as well as SNAT1 [53]) or “indirect,” possibly through its association with tetraspanins. Note, this is also true of CD81/other tetraspanins, which may likewise be “direct” or “indirect” targets (e.g. by their association with EWI-2). Our data do not distinguish these possibilities, and further mechanistic studies would be required to delineate the detailed mechanism of Vpu-mediated depletion. It should also be noted that in Table S1 of [64], EWI-2 depletion in CEM-T4 cells is (somewhat) dependent on the expression of Vpr. The effect size is modest and likely “indirect”, and does not contradict the Vpu and Nef data shown here. It does, however, suggest that the mechanism of EWI-2 depletion in HIV-1 infected T cells may be complex.

Overall, the combination of these two features (enrichment during assembly and transmission at the VS, and regulation by HIV-1 accessory proteins in infected cells), together with the fusion-preventing functions, strongly suggests that a particular host factor plays an important role in virus replication.

We expect that EWI-2 also inhibits the fusion of virus particles to target cells, as tetraspanins do [54,56,60], and we are currently testing that hypothesis (within the context of an extensive follow-up analysis aimed at dissecting the molecular determinants responsible for EWI-2’s fusion-inhibitory functions). It seems likely that tetraspanins and EWI-2 are not only tolerated but indeed enriched at virus budding sites because the benefit of cell-cell fusion inhibition at the VS is balanced against any negative effect of a reduction in virus infectivity. This is demonstrated by the fact that, in a native (unmanipulated) context, it is simultaneously true that (A) HIV-1-infected T cells routinely exhibit enrichment of these fusion inhibitors at virus release sites, (B) that cell-cell fusion is relatively infrequent, and (C) that HIV-1 spreads efficiently in those cell cultures.

As mentioned, while fusion inhibition operates at many levels and is orchestrated by HIV-1 proteins during infection, syncytia do nevertheless form, including *in vivo* [7–9] and when using a transmitted/founder (T/F) R5-tropic Env or even full-length replication-competent T/F virus [10,12]. However, these syncytia seem to remain small, at 4 or fewer nuclei and the vast majority having only 2 nuclei [9]. Very large syncytia (dozens to thousands of nuclei) are only induced by HIV-1 infection of certain T cell lines, especially Sup-T1 cells [65], or *in vivo* but only with the involvement of macrophage or dendritic cells [66–68]. It is therefore possible that T cell-T cell fusion is inhibited not only when a mononucleated infected cell encounters a target cell, but also when a syncytium encounters a target cell. An alternative explanation is that syncytia may be less viable as they grow larger, though some evidence contradicts that [69]. Here, we present evidence that host fusion-inhibitory proteins EWI-2 and CD81 are present at higher levels on the surface of small T cell syncytia when compared to mononucleated infected cells in the same culture. Because we find that the fusion-inhibitory capacity of EWI-2 and CD81 is also dose-dependent, it would therefore be expected that a higher “dose” of EWI-2 and/or CD81 in syncytia would make them less likely to undergo cell-cell fusion a second (or third) time. We are currently formally testing this hypothesis, and also investigating the surface levels on syncytia of other host proteins normally downregulated upon HIV-1 infection. Without implicating any particular fusion-inhibitory protein, we have in the past found evidence that indeed fusion-inhibitory factors may also be acting at syncytium-target cell VSs [9]: in Movie S7 of that report, we showed an example of a small syncytium containing 2 nuclei undergoing cell-cell fusion and acquiring a third nucleus. Subsequently, that syncytium encountered uninfected target cells and transferred virus particles to them through close contact, but did not undergo further cell-cell fusion and instead fully separated from them despite exhibiting the ability to fuse only hours earlier. We can now speculate that, as a result of the cell-cell fusion event we captured at the beginning of that sequence, this syncytium likely acquired a dose of EWI-2 and/or CD81, which subsequently allowed the syncytium to mediate cell-to-cell virus transfer at the VS without further cell-cell fusion.

Finally, repressing HIV-1 Env-induced cell-cell fusion not only allows for continued increase in the number of infected cells (as that number doubles each time producer and target cells separate after virus transmission), keeping Env’s fusion activity at bay may also be beneficial for the virus for other reasons. For instance, we and others have recently shown that lowering Env’s fusion activity also allows HIV-1 to overcome a restriction factor (APOBEC3G; [59]), and even antiviral drugs [70]. Further, large syncytia, that could form if Env-induced cell-cell fusion is uncontrolled, are likely prone to be attacked by innate immune cells. It is therefore critical that HIV-1 recruits fusion-inhibitory host factors such as EWI-2 to the VS to prevent excess cell-cell fusion and keep T cell syncytia small when they do form.

## Author Contributions

E.E.W., N.J.M., M.S., and M.T. conceived and designed the experiments. E.E.W. performed the experiments and analyzed the results, with contributions by M.S. and P.B.M. in Figure 1. N.J.M. performed the proteomics and analyzed the results in Figure 3A and Figure 4. S.P. performed the FG12 vector modification to remove the GFP reporter cassette. E.E.W., N.J.M., and M.S. prepared the figures. E.E.W., N.J.M., M.S., and M.T. wrote and edited the manuscript.

## Acknowledgments

The flow cytometry data we presented were obtained at the Harry Hood Bassett Flow Cytometry and Cell Sorting Facility, Larner College of Medicine, University of Vermont. The imaging shown in Figure 1A was performed at the Imaging/Physiology Core Facility, Neuroscience Center of Biomedical Research Excellence, Larner College of Medicine, University of Vermont.

## References

1. Jolly, C.; Kashefi, K.; Hollinshead, M.; Sattentau, Q.J. HIV-1 cell to cell transfer across an Env-induced, actin-dependent synapse. J Exp Med 2004, 199, 283–293, doi:10.1084/jem.20030648.

2. Hubner, W.; McNerney, G.P.; Chen, P.; Dale, B.M.; Gordon, R.E.; Chuang, F.Y.; Li, X.D.; Asmuth, D.M.; Huser, T.; Chen, B.K. Quantitative 3D video microscopy of HIV transfer across T cell virological synapses. Science 2009, 323, 1743–1747, doi:10.1126/science.1167525.

3. Ladinsky, M.S.; Kieffer, C.; Olson, G.; Deruaz, M.; Vrbanac, V.; Tager, A.M.; Kwon, D.S.; Bjorkman, P.J. Electron tomography of HIV-1 infection in gut-associated lymphoid tissue. PLoS Pathog 2014, 10, e1003899, doi:10.1371/journal.ppat.1003899.

4. Reh, L.; Magnus, C.; Schanz, M.; Weber, J.; Uhr, T.; Rusert, P.; Trkola, A. Capacity of Broadly Neutralizing Antibodies to Inhibit HIV-1 Cell-Cell Transmission Is Strain- and Epitope-Dependent. PLoS Pathog 2015, 11, e1004966, doi:10.1371/journal.ppat.1004966.

5. Bracq, L.; Xie, M.; Benichou, S.; Bouchet, J. Mechanisms for Cell-to-Cell Transmission of HIV-1. Front Immunol 2018, 9, 260, doi:10.3389/fimmu.2018.00260.

6. Imle, A.; Kumberger, P.; Schnellbacher, N.D.; Fehr, J.; Carrillo-Bustamante, P.; Ales, J.; Schmidt, P.; Ritter, C.; Godinez, W.J.; Muller, B., et al. Experimental and computational analyses reveal that environmental restrictions shape HIV-1 spread in 3D cultures. Nat Commun 2019, 10, 2144, doi:10.1038/s41467-019-09879-3.

7. Orenstein, J.M. In vivo cytolysis and fusion of human immunodeficiency virus type 1-infected lymphocytes in lymphoid tissue. J Infect Dis 2000, 182, 338–342, doi:10.1086/315640.

8. Murooka, T.T.; Deruaz, M.; Marangoni, F.; Vrbanac, V.D.; Seung, E.; von Andrian, U.H.; Tager, A.M.; Luster, A.D.; Mempel, T.R. HIV-infected T cells are migratory vehicles for viral dissemination. Nature 2012, 490, 283–287, doi:10.1038/nature11398.

9. Symeonides, M.; Murooka, T.T.; Bellfy, L.N.; Roy, N.H.; Mempel, T.R.; Thali, M. HIV-1-Induced Small T Cell Syncytia Can Transfer Virus Particles to Target Cells through Transient Contacts. Viruses 2015, 7, 6590–6603, doi:10.3390/v7122959.

10. Law, K.M.; Komarova, N.L.; Yewdall, A.W.; Lee, R.K.; Herrera, O.L.; Wodarz, D.; Chen, B.K. In Vivo HIV-1 Cell-to-Cell Transmission Promotes Multicopy Micro-compartmentalized Infection. Cell Rep 2016, 15, 2771–2783, doi:10.1016/j.celrep.2016.05.059.

11. Uchil, P.D.; Haugh, K.A.; Pi, R.; Mothes, W. In Vivo Imaging-Driven Approaches to Study Virus Dissemination and Pathogenesis. Annu Rev Virol 2019, 6, 501–524, doi:10.1146/annurev-virology-101416-041429.

12. Ventura, J.D.; Beloor, J.; Allen, E.; Zhang, T.; Haugh, K.A.; Uchil, P.D.; Ochsenbauer, C.; Kieffer, C.; Kumar, P.; Hope, T.J., et al. Longitudinal bioluminescent imaging of HIV-1 infection during antiretroviral therapy and treatment interruption in humanized mice. bioRxiv 2019, 10.1101/745125, 745125, doi:10.1101/745125.

13. Alvarez, R.A.; Barria, M.I.; Chen, B.K. Unique features of HIV-1 spread through T cell virological synapses. PLoS Pathog 2014, 10, e1004513, doi:10.1371/journal.ppat.1004513.

14. Compton, A.A.; Schwartz, O. They Might Be Giants: Does Syncytium Formation Sink or Spread HIV Infection? PLoS Pathog 2017, 13, e1006099, doi:10.1371/journal.ppat.1006099.

15. Rowell, J.F.; Stanhope, P.E.; Siliciano, R.F. Endocytosis of endogenously synthesized HIV-1 envelope protein. Mechanism and role in processing for association with class II MHC. J Immunol 1995, 155, 473–488.

16. Egan, M.A.; Carruth, L.M.; Rowell, J.F.; Yu, X.; Siliciano, R.F. Human immunodeficiency virus type 1 envelope protein endocytosis mediated by a highly conserved intrinsic internalization signal in the cytoplasmic domain of gp41 is suppressed in the presence of the Pr55gag precursor protein. J Virol 1996, 70, 6547–6556.

17. Roy, N.H.; Chan, J.; Lambele, M.; Thali, M. Clustering and mobility of HIV-1 Env at viral assembly sites predict its propensity to induce cell-cell fusion. J Virol 2013, 87, 7516–7525, doi:10.1128/jvi.00790-13.

18. Murakami, T.; Ablan, S.; Freed, E.O.; Tanaka, Y. Regulation of human immunodeficiency virus type 1 Env-mediated membrane fusion by viral protease activity. J Virol 2004, 78, 1026–1031.

19. Wyma, D.J.; Jiang, J.; Shi, J.; Zhou, J.; Lineberger, J.E.; Miller, M.D.; Aiken, C. Coupling of human immunodeficiency virus type 1 fusion to virion maturation: a novel role of the gp41 cytoplasmic tail. J Virol 2004, 78, 3429–3435.

20. Jiang, J.; Aiken, C. Maturation-dependent human immunodeficiency virus type 1 particle fusion requires a carboxyl-terminal region of the gp41 cytoplasmic tail. J Virol 2007, 81, 9999–10008, doi:10.1128/jvi.00592-07.

21. Chojnacki, J.; Staudt, T.; Glass, B.; Bingen, P.; Engelhardt, J.; Anders, M.; Schneider, J.; Muller, B.; Hell, S.W.; Krausslich, H.G. Maturation-dependent HIV-1 surface protein redistribution revealed by fluorescence nanoscopy. Science 2012, 338, 524–528, doi:10.1126/science.1226359.

22. Weng, J.; Krementsov, D.N.; Khurana, S.; Roy, N.H.; Thali, M. Formation of syncytia is repressed by tetraspanins in human immunodeficiency virus type 1-producing cells. J Virol 2009, 83, 7467–7474, doi:10.1128/jvi.00163-09.

23. Symeonides, M.; Lambele, M.; Roy, N.H.; Thali, M. Evidence showing that tetraspanins inhibit HIV-1-induced cell-cell fusion at a post-hemifusion stage. Viruses 2014, 6, 1078–1090, doi:10.3390/v6031078.

24. Roy, N.H.; Lambele, M.; Chan, J.; Symeonides, M.; Thali, M. Ezrin is a component of the HIV-1 virological presynapse and contributes to the inhibition of cell-cell fusion. J Virol 2014, 88, 7645–7658, doi:10.1128/jvi.00550-14.

25. Charrin, S.; Le Naour, F.; Oualid, M.; Billard, M.; Faure, G.; Hanash, S.M.; Boucheix, C.; Rubinstein, E. The major CD9 and CD81 molecular partner. Identification and characterization of the complexes. J Biol Chem 2001, 276, 14329–14337, doi:10.1074/jbc.M011297200.

26. Charrin, S.; Latil, M.; Soave, S.; Polesskaya, A.; Chretien, F.; Boucheix, C.; Rubinstein, E. Normal muscle regeneration requires tight control of muscle cell fusion by tetraspanins CD9 and CD81. Nat Commun 2013, 4, 1674, doi:10.1038/ncomms2675.

27. Sala-Valdes, M.; Ursa, A.; Charrin, S.; Rubinstein, E.; Hemler, M.E.; Sanchez-Madrid, F.; Yanez-Mo, M. EWI-2 and EWI-F link the tetraspanin web to the actin cytoskeleton through their direct association with ezrin-radixin-moesin proteins. J Biol Chem 2006, 281, 19665–19675, doi:10.1074/jbc.M602116200.

28. Stipp, C.S.; Kolesnikova, T.V.; Hemler, M.E. EWI-2 is a major CD9 and CD81 partner and member of a novel Ig protein subfamily. J Biol Chem 2001, 276, 40545–40554, doi:10.1074/jbc.M107338200.

29. Charrin, S.; Le Naour, F.; Labas, V.; Billard, M.; Le Caer, J.P.; Emile, J.F.; Petit, M.A.; Boucheix, C.; Rubinstein, E. EWI-2 is a new component of the tetraspanin web in hepatocytes and lymphoid cells. Biochem J 2003, 373, 409–421, doi:10.1042/bj20030343.

30. Rocha-Perugini, V.; Montpellier, C.; Delgrange, D.; Wychowski, C.; Helle, F.; Pillez, A.; Drobecq, H.; Le Naour, F.; Charrin, S.; Levy, S., et al. The CD81 partner EWI-2wint inhibits hepatitis C virus entry. PLoS One 2008, 3, e1866, doi:10.1371/journal.pone.0001866.

31. Montpellier, C.; Tews, B.A.; Poitrimole, J.; Rocha-Perugini, V.; D’Arienzo, V.; Potel, J.; Zhang, X.A.; Rubinstein, E.; Dubuisson, J.; Cocquerel, L. Interacting regions of CD81 and two of its partners, EWI-2 and EWI-2wint, and their effect on hepatitis C virus infection. J Biol Chem 2011, 286, 13954–13965, doi:10.1074/jbc.M111.220103.

32. Gordon-Alonso, M.; Sala-Valdes, M.; Rocha-Perugini, V.; Perez-Hernandez, D.; Lopez-Martin, S.; Ursa, A.; Alvarez, S.; Kolesnikova, T.V.; Vazquez, J.; Sanchez-Madrid, F., et al. EWI-2 association with alpha-actinin regulates T cell immune synapses and HIV viral infection. J Immunol 2012, 189, 689–700, doi:10.4049/jimmunol.1103708.

33. Chen, A.; Leikina, E.; Melikov, K.; Podbilewicz, B.; Kozlov, M.M.; Chernomordik, L.V. Fusion-pore expansion during syncytium formation is restricted by an actin network. J Cell Sci 2008, 121, 3619–3628, doi:10.1242/jcs.032169.

34. Maddon, P.J.; Dalgleish, A.G.; Mcdougal, J.S.; Clapham, P.R.; Weiss, R.A.; Axel, R. The T4 Gene Encodes the Aids Virus Receptor and Is Expressed in the Immune-System and the Brain. Cell 1986, 47, 333–348, doi:Doi 10.1016/0092-8674(86)90590-8.

35. Platt, E.J.; Wehrly, K.; Kuhmann, S.E.; Chesebro, B.; Kabat, D. Effects of CCR5 and CD4 cell surface concentrations on infections by macrophagetropic isolates of human immunodeficiency virus type 1. J Virol 1998, 72, 2855–2864.

36. Derdeyn, C.A.; Decker, J.M.; Sfakianos, J.N.; Wu, X.; O’Brien, W.A.; Ratner, L.; Kappes, J.C.; Shaw, G.M.; Hunter, E. Sensitivity of human immunodeficiency virus type 1 to the fusion inhibitor T-20 is modulated by coreceptor specificity defined by the V3 loop of gp120. J Virol 2000, 74, 8358–8367.

37. Wei, X.; Decker, J.M.; Liu, H.; Zhang, Z.; Arani, R.B.; Kilby, J.M.; Saag, M.S.; Wu, X.; Shaw, G.M.; Kappes, J.C. Emergence of resistant human immunodeficiency virus type 1 in patients receiving fusion inhibitor (T-20) monotherapy. Antimicrob. Agents Chemother. 2002, 46, 1896–1905.

38. Takeuchi, Y.; McClure, M.O.; Pizzato, M. Identification of gammaretroviruses constitutively released from cell lines used for human immunodeficiency virus research. J Virol 2008, 82, 12585–12588, doi:10.1128/jvi.01726-08.

39. Platt, E.J.; Bilska, M.; Kozak, S.L.; Kabat, D.; Montefiori, D.C. Evidence that ecotropic murine leukemia virus contamination in TZM-bl cells does not affect the outcome of neutralizing antibody assays with human immunodeficiency virus type 1. J Virol 2009, 83, 8289–8292, doi:10.1128/JVI.00709-09.

40. Trkola, A.; Matthews, J.; Gordon, C.; Ketas, T.; Moore, J.P. A cell line-based neutralization assay for primary human immunodeficiency virus type 1 isolates that use either the CCR5 or the CXCR4 coreceptor. J Virol 1999, 73, 8966–8974.

41. Spenlehauer, C.; Gordon, C.A.; Trkola, A.; Moore, J.P. A luciferase-reporter gene-expressing T-cell line facilitates neutralization and drug-sensitivity assays that use either R5 or X4 strains of human immunodeficiency virus type 1. Virology 2001, 280, 292–300, doi:10.1006/viro.2000.0780.

42. Foley, G.E.; Lazarus, H.; Farber, S.; Uzman, B.G.; Boone, B.A.; McCarthy, R.E. Continuous Culture of Human Lymphoblasts from Peripheral Blood of a Child with Acute Leukemia. Cancer 1965, 18, 522–529.

43. Nara, P.L.; Hatch, W.C.; Dunlop, N.M.; Robey, W.G.; Arthur, L.O.; Gonda, M.A.; Fischinger, P.J. Simple, rapid, quantitative, syncytium-forming microassay for the detection of human immunodeficiency virus neutralizing antibody. AIDS Res Hum Retroviruses 1987, 3, 283–302, doi:10.1089/aid.1987.3.283.

44. Nara, P.L.; Fischinger, P.J. Quantitative infectivity assay for HIV-1 and-2. Nature 1988, 332, 469–470, doi:10.1038/332469a0.

45. Refsland, E.W.; Hultquist, J.F.; Harris, R.S. Endogenous origins of HIV-1 G-to-A hypermutation and restriction in the nonpermissive T cell line CEM2n. PLoS Pathog 2012, 8, e1002800, doi:10.1371/journal.ppat.1002800.

46. Simm, M.; Shahabuddin, M.; Chao, W.; Allan, J.S.; Volsky, D.J. Aberrant Gag protein composition of a human immunodeficiency virus type 1 vif mutant produced in primary lymphocytes. J Virol 1995, 69, 4582–4586.

47. Freed, E.O.; Martin, M.A. Virion incorporation of envelope glycoproteins with long but not short cytoplasmic tails is blocked by specific, single amino acid substitutions in the human immunodeficiency virus type 1 matrix. J Virol 1995, 69, 1984–1989.

48. Qin, X.F.; An, D.S.; Chen, I.S.; Baltimore, D. Inhibiting HIV-1 infection in human T cells by lentiviral-mediated delivery of small interfering RNA against CCR5. Proc Natl Acad Sci U S A 2003, 100, 183–188, doi:10.1073/pnas.232688199.

49. Schindelin, J.; Arganda-Carreras, I.; Frise, E.; Kaynig, V.; Longair, M.; Pietzsch, T.; Preibisch, S.; Rueden, C.; Saalfeld, S.; Schmid, B., et al. Fiji: an open-source platform for biological-image analysis. Nat Methods 2012, 9, 676–682, doi:10.1038/nmeth.2019.

50. Greenwood, E.J.; Matheson, N.J.; Wals, K.; van den Boomen, D.J.; Antrobus, R.; Williamson, J.C.; Lehner, P.J. Temporal proteomic analysis of HIV infection reveals remodelling of the host phosphoproteome by lentiviral Vif variants. Elife 2016, 5, doi:10.7554/eLife.18296.

51. Naamati, A.; Williamson, J.C.; Greenwood, E.J.; Marelli, S.; Lehner, P.J.; Matheson, N.J. Functional proteomic atlas of HIV infection in primary human CD4+ T cells. Elife 2019, 8, doi:10.7554/eLife.41431.

52. Matheson, N.J.; Peden, A.A.; Lehner, P.J. Antibody-free magnetic cell sorting of genetically modified primary human CD4+ T cells by one-step streptavidin affinity purification. PLoS One 2014, 9, e111437, doi:10.1371/journal.pone.0111437.

53. Matheson, N.J.; Sumner, J.; Wals, K.; Rapiteanu, R.; Weekes, M.P.; Vigan, R.; Weinelt, J.; Schindler, M.; Antrobus, R.; Costa, A.S., et al. Cell Surface Proteomic Map of HIV Infection Reveals Antagonism of Amino Acid Metabolism by Vpu and Nef. Cell Host Microbe 2015, 18, 409–423, doi:10.1016/j.chom.2015.09.003.

54. Krementsov, D.N.; Weng, J.; Lambele, M.; Roy, N.H.; Thali, M. Tetraspanins regulate cell-to-cell transmission of HIV-1. Retrovirology 2009, 6, 64, doi:10.1186/1742-4690-6-64.

55. Durham, N.D.; Chen, B.K. HIV-1 Cell-Free and Cell-to-Cell Infections Are Differentially Regulated by Distinct Determinants in the Env gp41 Cytoplasmic Tail. J Virol 2015, 89, 9324–9337, doi:10.1128/JVI.00655-15.

56. Lambele, M.; Koppensteiner, H.; Symeonides, M.; Roy, N.H.; Chan, J.; Schindler, M.; Thali, M. Vpu is the main determinant for tetraspanin downregulation in HIV-1-infected cells. J Virol 2015, 89, 3247–3255, doi:10.1128/jvi.03719-14.

57. Haller, C.; Muller, B.; Fritz, J.V.; Lamas-Murua, M.; Stolp, B.; Pujol, F.M.; Keppler, O.T.; Fackler, O.T. HIV-1 Nef and Vpu are functionally redundant broad-spectrum modulators of cell surface receptors, including tetraspanins. J Virol 2014, 88, 14241–14257, doi:10.1128/jvi.02333-14.

58. Karn, J.; Stoltzfus, C.M. Transcriptional and posttranscriptional regulation of HIV-1 gene expression. Cold Spring Harb Perspect Med 2012, 2, a006916, doi:10.1101/cshperspect.a006916.

59. Ikeda, T.; Symeonides, M.; Albin, J.S.; Li, M.; Thali, M.; Harris, R.S. HIV-1 adaptation studies reveal a novel Env-mediated homeostasis mechanism for evading lethal hypermutation by APOBEC3G. PLoS Pathog 2018, 14, e1007010, doi:10.1371/journal.ppat.1007010.

60. Sato, K.; Aoki, J.; Misawa, N.; Daikoku, E.; Sano, K.; Tanaka, Y.; Koyanagi, Y. Modulation of human immunodeficiency virus type 1 infectivity through incorporation of tetraspanin proteins. J Virol 2008, 82, 1021–1033, doi:10.1128/jvi.01044-07.

61. Sugden, S.M.; Bego, M.G.; Pham, T.N.; Cohen, E.A. Remodeling of the Host Cell Plasma Membrane by HIV-1 Nef and Vpu: A Strategy to Ensure Viral Fitness and Persistence. Viruses 2016, 8, 67, doi:10.3390/v8030067.

62. Gordon-Alonso, M.; Yanez-Mo, M.; Barreiro, O.; Alvarez, S.; Munoz-Fernandez, M.A.; Valenzuela-Fernandez, A.; Sanchez-Madrid, F. Tetraspanins CD9 and CD81 modulate HIV-1-induced membrane fusion. J Immunol 2006, 177, 5129–5137, doi:10.4049/jimmunol.177.8.5129.

63. Len, A.C.L.; Starling, S.; Shivkumar, M.; Jolly, C. HIV-1 Activates T Cell Signaling Independently of Antigen to Drive Viral Spread. Cell Rep 2017, 18, 1062–1074, doi:10.1016/j.celrep.2016.12.057.

64. Greenwood, E.J.D.; Williamson, J.C.; Sienkiewicz, A.; Naamati, A.; Matheson, N.J.; Lehner, P.J. Promiscuous Targeting of Cellular Proteins by Vpr Drives Systems-Level Proteomic Remodeling in HIV-1 Infection. Cell Rep 2019, 27, 1579–1596 e1577, doi:10.1016/j.celrep.2019.04.025.

65. Sylwester, A.; Murphy, S.; Shutt, D.; Soll, D.R. HIV-induced T cell syncytia are self-perpetuating and the primary cause of T cell death in culture. J Immunol 1997, 158, 3996–4007.

66. Rinfret, A.; Latendresse, H.; Lefebvre, R.; St-Louis, G.; Jolicoeur, P.; Lamarre, L. Human immunodeficiency virus-infected multinucleated histiocytes in oropharyngeal lymphoid tissues from two asymptomatic patients. Am J Pathol 1991, 138, 421–426.

67. Frankel, S.S.; Wenig, B.M.; Burke, A.P.; Mannan, P.; Thompson, L.D.; Abbondanzo, S.L.; Nelson, A.M.; Pope, M.; Steinman, R.M. Replication of HIV-1 in dendritic cell-derived syncytia at the mucosal surface of the adenoid. Science 1996, 272, 115–117, doi:10.1126/science.272.5258.115.

68. Murooka, T.T.; Sharaf, R.R.; Mempel, T.R. Large Syncytia in Lymph Nodes Induced by CCR5-Tropic HIV-1. AIDS Res Hum Retroviruses 2015, 31, 471–472, doi:10.1089/aid.2014.0378.

69. Sylwester, A.; Daniels, K.; Soll, D.R. The invasive and destructive behavior of HIV-induced T cell syncytia on collagen and endothelium. J Leukoc Biol 1998, 63, 233–244, doi:10.1002/jlb.63.2.233.

70. Van Duyne, R.; Kuo, L.S.; Pham, P.; Fujii, K.; Freed, E.O. Mutations in the HIV-1 envelope glycoprotein can broadly rescue blocks at multiple steps in the virus replication cycle. Proc Natl Acad Sci U S A 2019, 116, 9040–9049, doi:10.1073/pnas.1820333116.

